# Population-scale single-cell RNA-seq profiling across dopaminergic neuron differentiation

**DOI:** 10.1101/2020.05.21.103820

**Authors:** J Jerber, DD Seaton, ASE Cuomo, N Kumasaka, J Haldane, J Steer, M Patel, D Pearce, M Andersson, MJ Bonder, E Mountjoy, M Ghoussaini, MA Lancaster, HipSci Consortium, JC Marioni, FT Merkle, O Stegle, DJ Gaffney

## Abstract

Common genetic variants can have profound effects on cellular function, but studying these effects in primary human tissue samples and during development is challenging. Human induced pluripotent stem cell (iPSC) technology holds great promise for assessing these effects across different differentiation contexts. Here, we use an efficient pooling strategy to differentiate 215 iPS cell lines towards a midbrain neural fate, including dopaminergic neurons, and profile over 1 million cells sampled across three differentiation timepoints using single cell RNA sequencing. We find that the proportion of neuronal cells produced by each cell line is highly reproducible over different experimental batches, and identify robust molecular markers in pluripotent cells that predict line-to-line differences in cell fate. We identify expression quantitative trait loci (eQTL) that manifest at different stages of neuronal development, and in response to oxidative stress, by exposing cells to rotenone. We find over one thousand eQTL that colocalise with a known risk locus for a neurological trait, nearly half of which are not found in GTEx. Our study illustrates how coupling single cell transcriptomics with long-term iPSC differentiation can profile mechanistic effects of human trait-associated genetic variants in otherwise inaccessible cell states.

## Introduction

Genetic variation can significantly alter cell function, for example by altering gene expression. Human Induced Pluripotent Stem Cells (iPSCs) are a promising cellular model for assessing the cellular consequences of human genetic variation across different lineages, developmental states and cell types. In particular, human iPSCs facilitate the study of developmental time points and stimulation conditions that would be challenging to obtain *in vivo*. The creation of cell banks containing hundreds of iPSC lines^1^ provides an exciting opportunity to carry out population-scale studies *in vitr*^2–5^. However, differentiating iPSCs is expensive and labour-intensive, and differentiation experiments are difficult to compare due to substantial batch variation. Thus, studies of more than a handful of lines remain a significant challenge. Furthermore, most iPSC differentiation protocols produce a heterogenous population of cells of which the target cell type is a subset^6–8^. This variability in differentiation outcomes hinders efforts to dissect the genetic contributions to cellular phenotypes.

Single cell sequencing has enabled “multiplexed” experimental designs, where cells from multiple donors are pooled together^2,9^. Pooling improves throughput and allows experimental variability between differentiation batches to be rigorously controlled, by enabling cell type heterogeneity to be accounted for in downstream analysis. To date, multiplexed experimental designs have only been applied to short differentiation protocols (over a period of days), that generate cells corresponding to very early stages of development, and have not captured developmental progression toward a mature cell fate. Population-scale pooling during long-term differentiation offers the opportunity to examine the effect of common genetic variants on gene expression in each cell population produced over neural development, providing a foundation for future mechanistic studies.

Here, we develop and apply a multiplexing strategy to profile the differentiation and maturation of more than two hundred iPSC lines derived from the Human Induced Pluripotent Stem Cell Initiative (HipSci) towards a midbrain neural fate, including dopaminergic neurons (DA). DA are involved in motor function and other cognitive processes and play key roles in neurological disorders, including Parkinson’s Disease^10,11^ (PD). To study how these cells differentiate, and how genetic background could influence differentiation, we employed a well-established protocol^12^ and collected cells at three maturation stages (progenitor-like, young neurons, and more mature neurons), covering 52 days of differentiation. We additionally exposed cells on day 51 to rotenone, to explore how genetic variation shapes the neuronal response to oxidative stress. Using this system, we create the first map of expression quantitative trait loci (eQTL) at multiple stages of human neuronal differentiation, and identify nearly 500 novel trait / eQTL colocalisations. Using estimates of cell population composition based on single cell RNA-seq, we demonstrate that a strong, cell intrinsic-differentiation bias affects a significant proportion of iPSC lines, such that approximately 25% reproducibly fail to produce any neuronal cells.

## Results

### High-throughput differentiation of midbrain dopaminergic neurons

We selected 215 iPSC lines derived from healthy donors by the HipSci project^1^ for differentiation towards a midbrain cell fate, including dopaminergic neurons^12^. Differentiation experiments were multiplexed using pools containing between 7 and 24 cell lines per experiment (**Supplementary Table 1**). Immunochemistry confirmed that cells from both pooled and conventional differentiation of individual lines expressed protein markers associated with patterning of DA (LMX1A, FOXA2 and TH) (**Supplementary Fig. 1**). To capture transcriptional changes during neurogenesis and neuronal maturation, we performed single cell RNA sequencing (scRNA-seq) of cells captured at day 11 (D11, midbrain floorplate progenitors), day 30 (D30, young post-mitotic midbrain neurons) and day 52 (D52, more mature midbrain neurons). To mimic an oxidative stress condition, we also profiled day 52 neurons upon exposure to a sub-lethal dose of rotenone (ROT, 0.1 μM; 24 h) a chemical stressor that preferentially leads to DA death in models of PD (**Fig. 1a**)^13^.

**Figure 1.**
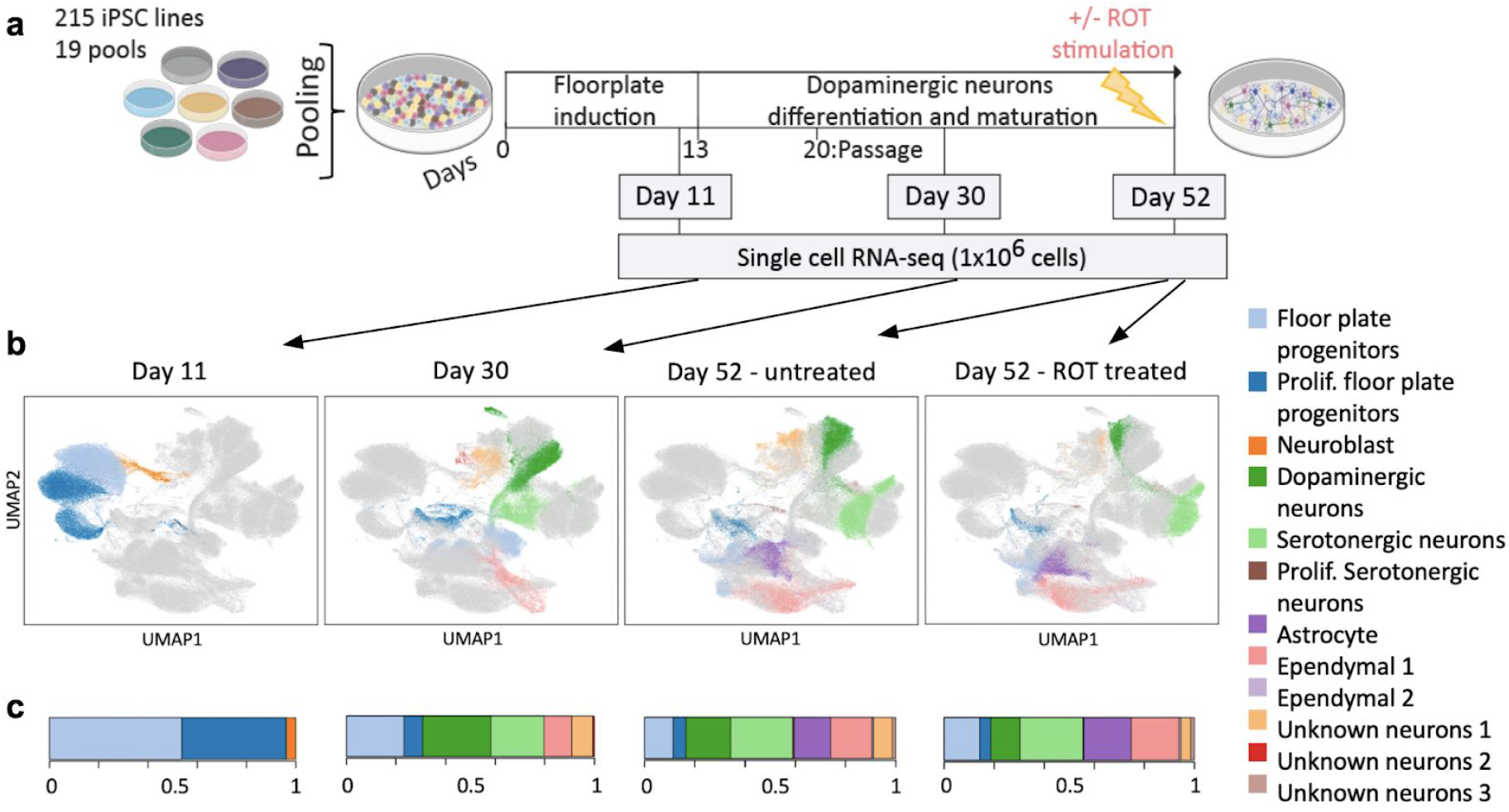
Experimental design and cell type heterogeneity in pooled differentiations of iPSCs to a midbrain cell fate. (a) Experimental workflow for scRNA-seq analysis of iPSC-derived dopaminergic neurons. Time points at which cells were collected for scRNA-seq profiling (Day 11, Day 30, Day 52) are indicated. On day 51, half of the cells were stimulated with rotenone (ROT) for 24h to induce oxidative stress. (b) UMAP plot of all 1,027,401 cells assayed, coloured by annotated cell type identity. Cells that were not collected at a given condition (time point, stimulus) are displayed in light grey. Prolif: Proliferating. (c) Barplot showing, for each condition, the fraction of cells assigned to each cell type.

After QC, we obtained a total of 1,027,401 cells across 19 cell pools^14^ and four conditions (**Fig. 1a, Supplementary Table 1**). The cell line of origin for each cell was inferred from single cell RNA-seq read data using known genotypes made available by the HipSci consortium (using demuxlet,^15^). Adjustment for experimental batch effects using Harmony^16^ followed by Louvain clustering^14^ identified a total of 26 clusters (6, 7 and 13 clusters respectively at D11, D30, D52, **Supplementary Fig. 2a**). These clusters were then assigned to cell types by testing for enrichments of 48 literature-curated marker genes of major brain cell types (**Fig. 1b, Supplementary Fig. 2b,c; Methods**).

Among these, we identified six dominant cell types that were making up at least 10% of the cells at any time point (**Supplementary Fig. 2c**). These included two main cell type populations at day 11: proliferating and non-proliferating midbrain floorplate progenitors (both expressing *LMX1A, FOXA2* and expressing *MIK67, TOP2A* when proliferating^17^). At days 30 and 52, four additional dominant cell types were identified. Two of these additional cell types appeared neuronal and two were non-neuronal (characterised by expression and lack of the pan-neuronal markers *SNAP25* and *SYT1* respectively). The two neuronal populations could be further divided into midbrain dopaminergic neurons, which expressed *NR4A2, PBX1, TMCC3*^17–19^ and serotonergic neurons (Sert), which expressed *TPH1, TPH2, GATA2*^20^. The two non-neuronal cell types corresponded to ependymal-like cells, detected both at days 30 and 52 (Ependymal 1^21^) and astrocyte-like cells, unique to day 52 (Astrocyte-like^22,23^). We also identified a neuroblast population, specific to day 11 (4% of all cells at day 11) expressing pro-neuronal genes (*NEUROD1, NEUROG2, NHLH1*^24,25^) and an additional neuronal population (expressing *SNAP25* and *SYT1*) that could not be assigned to a specific neuronal identity (Unknown neurons 1, present at Day 30 and Day 52 at around 7%). Finally, we identified four additional rare cell types (<2% of cells sampled at any time point), including a second ependymal-like population (Ependymal 2), a proliferating neuronal serotonergic population (Prolif. Serotonergic neurons), and two additional neuronal populations which could not be annotated unambiguously (Unknown neurons 2, Unknown neurons 3, **Fig. 1b, Supplementary Fig. 2c**).

UMAP projection of cells collected across all time points, stimuli and lines revealed broad co-clustering of cell types, but with noticeable differences between time points and stimuli (**Fig. 1b**). We observed substantial variation in the cell type proportions across time points and stimuli (**Fig. 1c**). For example, the proportion of DA upon ROT stimulation was significantly reduced (30% reduction upon stimulation, Fisher’s exact test, p=2.2×10^−16^), consistent with previous observations that dopaminergic neurons are most affected by apoptosis due to oxidative stress^26–28^.

Collectively, our population-scale scRNA-seq analysis revealed a surprisingly diverse repertoire of cell types, enabling study of both cell line differentiation propensity and the identification of genetic variants that act in a cell type-specific manner.

### Intrinsic variation in differentiation efficiency between iPSC lines

iPSC differentiation protocols are highly variable but the biological basis for this remains largely obscure, which hinders efforts to rationally select cell lines for specific applications^29,30^. We found substantial variation in the proportions of different cell types produced by different iPSC cell lines at each time point (**Fig. 2a**, **Supplementary Fig. 3a, Supplementary Table 2**). Using principal component analysis of cell type fractions per cell line and pool, we identified the proportion of midbrain neurons (DA and Sert) on day 52 as the largest axis of variation (PC1, 47% variance, **Supplementary Fig. 3b-d**). Since DA and Sert cells are derived from similar progenitor populations *in vivo*, it is not surprising that both populations are observed in our differentiation experiment^31^. This motivated us to estimate “differentiation efficiency” for each iPSC line, defined as the sum of the proportions of DA and Sert cells produced on day 52 (**Fig. 2b**). We assessed the reproducibility of this measure of differentiation efficiency using data from 35 lines that were differentiated twice, in two different pools. Importantly, we found that iPSC line differentiation efficiency defined in this way was highly reproducible between different pools (Pearson R=0.75; p=2×10^−6^; **Fig. 2d**).

**Figure 2.**
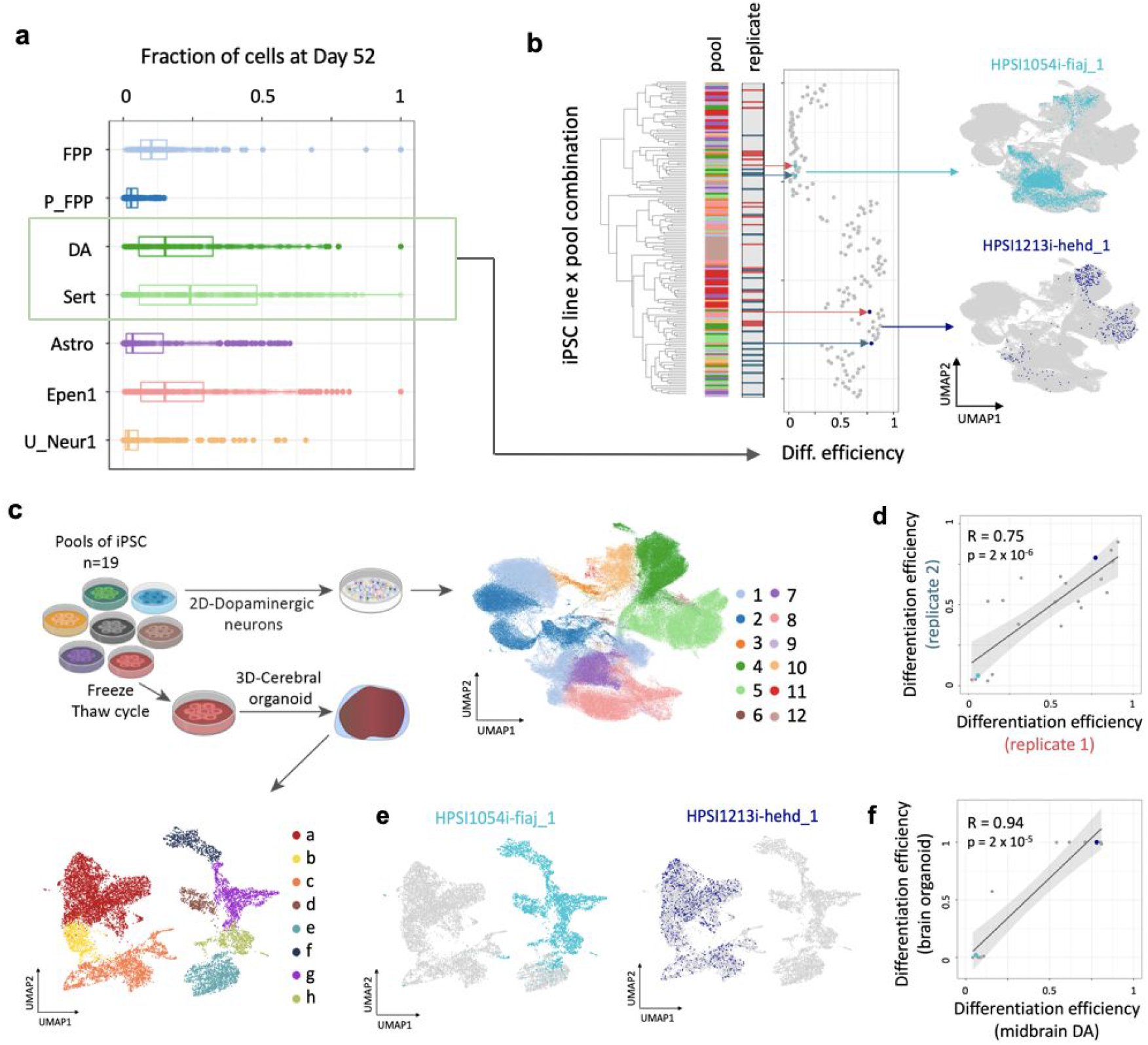
Reproducible variation in differentiation trajectories. (a) Box plot showing, for each cell type, the proportions of that cell type across cell lines on day 52. Each point indicates a different cell line. We defined the sum of the proportions of DA and Sert cells generated at day 52 as differentiation efficiency. (b) Hierarchical clustering of (cell line,pool) combinations by differentiation efficiency. Pools are shown in the first bar and the colours indicate in which of 10 pools (for which we had data at all time points, used to define differentiation efficiency; Supplementary Fig. 3, Methods) each line was differentiated. Differentiation replicates, where the same line was present in 2 pools, are shown in the second bar (replicate 1 in red and replicate 2 in blue). UMAPs, highlighting the distributions of cells on day 52 for two selected cell lines with low and high differentiation efficiencies respectively (HPSI0514i-fiaj_1, in seagreen and HPSI1213i-hehd_1, in dark blue) (right). (c) Experimental workflow for scRNA-seq analysis of iPSC-derived cerebral organoids using one pool consisting of 18 cell lines, profiled using scRNA-seq after 113 days of differentiation. UMAPs of the two resulting cell populations. UMAP legend: 1: FPP, 2: P_FPP 3: Neuroblasts, 4: DA, 5: Sert, 6: Proliferating Sert, 7: Astro, 8: Epen1, 9: Epend2, 10: U_Neur1, 11: U_Neur2, 12: U_Neur3; a: Neural cells, b: Intermediate progenitors, c: Radial glial progenitors, d: Satellite cells, e: Mesenchymal cells, f: Myotube, g: PAX7+ cells, h: Wnt+cells. (d) Scatter plot showing estimated differentiation efficiency between differentiation replicates (i.e. cell lines differentiated in two different pools, n=35). Highlighted are the two cell lines from (b). (e) UMAPs of two representative cell lines making non-brain and brain cell types in the organoid study. The two cell lines selected are the same as in b. (f) Scatter plot of differentiation efficiency as measured using midbrain dopaminergic neuronal differentiation (x-axis) versus differentiation efficiency as measured in organoid differentiation (y-axis) for a subset of 12 iPS cell lines in common. Highlighted are the two cell lines from (b). Astro: Astrocyte-like; DA: Dopaminergic neurons, Epen1: Ependymal-like1, FPP: Floor Plate Progenitors, P_FPP: Proliferating FPP, Sert: Serotonergic neurons, U_Neur1,2,3: Unknown neurons 1,2,3.

Given the robustness of these results, we wondered if they were generalisable to other neuronal differentiation approaches. We therefore differentiated a pool of 18 lines (pool 4) into cerebral organoids for 113 days^32^ and profiled the resulting cell populations using scRNA-seq (11,445 cells, **Fig. 2c, Methods**). Remarkably, we found that the proportion of brain cell types (all neural, glial, and neural progenitor cells, **Supplementary Fig. 4**) produced by each line in the cerebral organoids was strongly correlated with differentiation efficiency as estimated from the dopaminergic differentiation (**Fig. 2e,f**, R=0.94; p=2×10^−5^; n=12). Taken together, these results strongly suggest that variation in iPSC neural differentiation efficiencies arise primarily due to cell-intrinsic factors. Furthermore, the consistency of differentiation efficiency suggests these properties extend to neuronal differentiation more generally.

### iPSC gene expression signatures predict neuronal differentiation efficiency

Motivated by the reproducibility of differentiation outcomes across multiple independent pools, we tested for associations between differentiation efficiency and other experimental and biological factors (**Supplementary Table 3)**, but found no or weak associations with cell line passage number (p=0.77), donor sex (p=0.008), chromosome X activation status (p=0.01, **Methods**), or PluriTest scores33 (p=0.01, **Methods**).

Next, we assessed whether differentiation efficiency was associated with particular patterns of gene expression in undifferentiated iPSCs. Using data from independent bulk RNA-seq data available for a subset of 184 iPSC lines included in this study^1,34^ we identified significant associations with differentiation efficiency for 2,045 genes (983 positive and 1,062 negative associations; F-test, FDR < 5%; **Fig. 3b,c; Supplementary Table 4, Methods**). Notably, when defining poor differentiation as a binary outcome (differentiation efficiency < 0.2), genome-wide gene expression profiles in undifferentiated iPSCs were able to predict differentiation outcomes (logistic regression; 100% precision at 35% recall assessed using cross-validation; **Supplementary Fig. 5a,b; Methods**). This result was robust to alternative thresholds for defining poor differentiation outcomes. Using this model, we generated predicted scores for all 812 HipSci lines with bulk RNA-seq data (**Supplementary Table 5**). This analysis indicated that a substantial fraction of lines in the HipSci resource (26%) were predicted to produce < 20% neuronal cells under the differentiation conditions we tested. Furthermore, we tested whether the same experimental and biological factors previously associated with differentiation efficiency replicated in this larger sample and found consistent results. (**Supplementary Table 3; Methods**). Finally, we also observed substantial variation in predicted outcomes of different lines from the same donor (**Supplementary Fig. 5c-e**), suggesting that donor genetic background is unlikely to play a role in driving differentiation biases.

**Figure 3.**
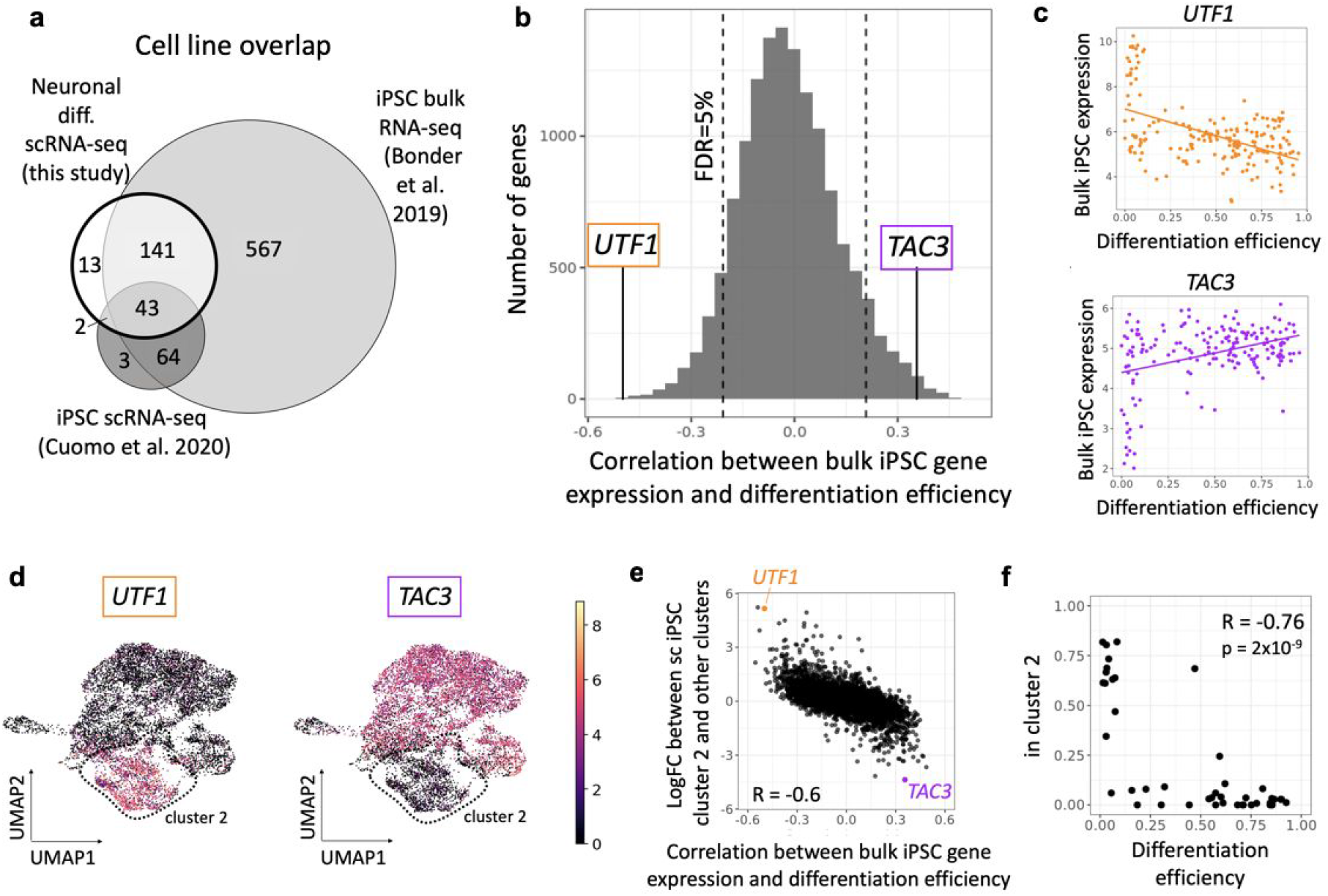
A gene expression signature in iPSCs is associated with neuronal differentiation efficiency. (a) Venn diagram indicating the overlap of cell lines included in this study and two recent iPSC studies, a bulk RNA-seq study^34^ and a single cell RNA-seq study^2^. (b) Histogram of Pearson correlation coefficients between variation in gene expression of individual genes (from bulk RNA-seq^34^) and differentiation efficiency. Two representative genes (*UTF1, TAC3*) are highlighted. (c) Example of genes from (b) whose gene expression in iPSC (based on bulk RNA-seq) is negatively (*UTF1*) and positively (*TAC3*) correlated with differentiation efficiency. (d) UMAPs of single-cell RNA-seq profiles in iPSCs from 112 donors^2^. Colours denote the expression level of the two example genes from (b),(c): *UTF1* and *TAC3*. Cluster 2 is shown by the dashed lines. (e) Comparison of marker gene association results with expression markers of the cluster 2. For each gene, the Pearson correlation coefficient of association between the gene and differentiation efficiency (x-axis; iPSC gene expression assessed using bulk RNA-seq, as in (b)) is compared to its log fold change between cluster 2 and the other clusters (y-axis, scRNA-seq). (*UTF1, TAC3*) are highlighted. (f) Scatterplot between the proportion of cells assigned to cluster 2 (y-axis) and differentiation efficiency (x-axis) across 45 cell lines which were included in both sets of experiments. Where multiple measurements were available for a given cell line, these were averaged.

Next, since iPSC cultures are heterogeneous, we hypothesised that the identified predictive gene signatures might arise from varying proportions of an iPSC subpopulation. To test this hypothesis, we re-analysed scRNA-seq data from 112 iPSC lines that were assayed previously under iPSC culture conditions similar to those used here^2^, 45 of which were also included in this study, **Methods**, **Fig. 3a**). We identified 5 clusters, all but one of which expressed similarly high levels of core pluripotency markers (*NANOG, SOX2, POU5F1*, **Supplementary Fig. 6a,b; Methods**). We found that genes whose expression predicted poor differentiation (e.g *UTF1*) were highly enriched in one of those clusters (cluster 2), while genes whose expression were predictive of successful differentiation (*TAC3*), were downregulated in cluster 2 relative to the remaining iPSC clusters (**Fig. 3d,e**). In comparison, other cell clusters did not show such equivalent enrichment in differentiation marker genes (**Supplementary Table 6**).

As a direct confirmation of this hypothesis, we also tested for and confirmed an association between the fraction of cells in cluster 2 and differentiation efficiency for each cell line (Pearson R=-0.76, p=2.05 x10^−9^; **Fig. 3f**). We used additional data from 2 to assess the consistency of the fraction of cluster 2 cells across replication experiments, finding high concordance (Pearson R=0.9; n=23, **Supplementary Fig. 6c**). Using the known relationship between iPSC bulk RNA-seq and the proportion of cluster 2 cells, we predicted this proportion for 182 cell lines included in our differentiation experiments, and confirmed the negative correlation with differentiation efficiency (Pearson R=-0.49; p=3×10^−12^, **Supplementary Fig. 6d; Methods**). Finally, we also analysed an additional scRNA-seq dataset from iPSCs derived from Lymphoblastoid Cell Lines^35^ (LCLs). Using our single cell analysis pipeline, we also identified a cluster of cells with a concordant expression profile to cluster 2 (**Methods, Supplementary Fig. 7**). Taken together, these results provide further evidence that a subpopulation of iPS cells with poor differentiation capability is consistently detected across different human iPSCs banks, and that this bias can be robustly predicted using expression markers at the iPSC stage. Importantly, despite the variability in differentiation efficiency, our single-cell sequencing approach enabled us to examine multiple disease-relevant cell populations for many cell lines. Correlating changes in gene expression with common genetic variants associated with various traits in GWAS could provide insights into disease mechanisms.

### eQTL discovery and comparison with *in vivo* eQTL maps

We next focused on understanding how individual-to-individual genetic variation influenced gene expression across these cell types during differentiation and in response to stimulation. Specifically, we mapped *cis* expression quantitative trait loci (eQTL) separately for each of the 14 distinct cell populations that corresponds to the profiled “cell type”-“condition” contexts. eQTL were mapped by calculating aggregate expression levels for each donor, considering common gene-proximal variants (MAF>0.05, plus or minus 250 kb around genes; **Methods**). Variability in differentiation efficiency between lines resulted in substantial differences in the number of cells collected for each donor (**Supplementary Fig. 8a**), affecting accuracy of the estimates of aggregated expression. To account for this source of noise, we adapted commonly used eQTL mapping strategies^2^ based on linear mixed models (LMMs) by incorporating an additional variance component into the model (**Methods**). This approach greatly increased the power to map eQTL, resulting in a total of 4,087 genes with at least one eQTL in any of the contexts (hereafter “eGene”, FDR < 5%, **Fig. 4a, Supplementary Fig. 8b, Supplementary Table 7**).

**Figure 4.**
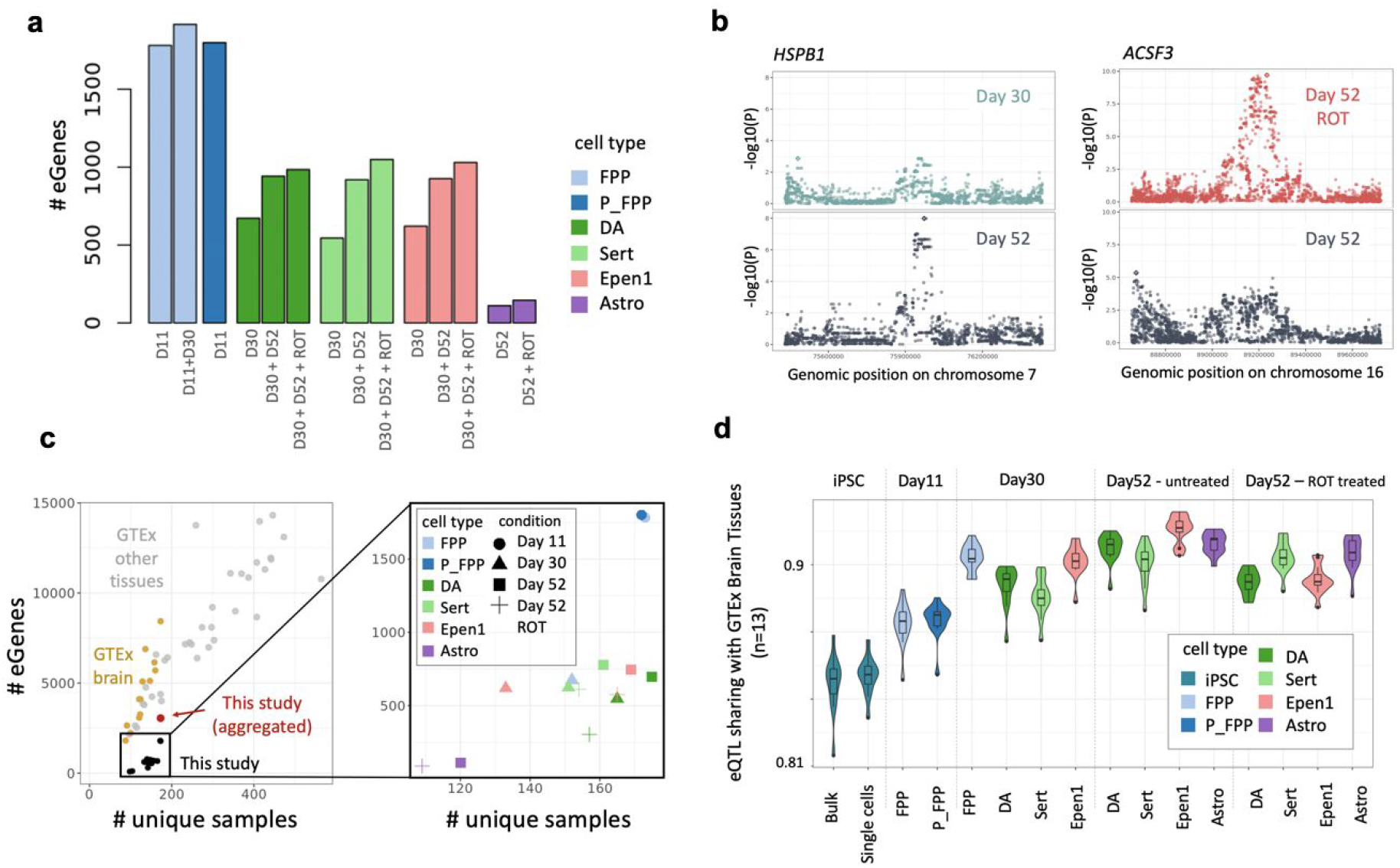
Mapping of *cis* eQTL in distinct cell contexts (“cell types”-“conditions”) identified across midbrain differentiation. (a) Cumulative number of genes with at least one eQTL (eGenes) for each cell type and condition (D11 = Day 11; D30 = Day 30; D52 = Day 52; ROT = Rotenone stimulation). (b) Left: day 52-specific eQTL for *HSPB1* in DA (rs6465098; FDR < 5%, Methods). Shown are Manhattan plots for DA at day 30 (top), day 52 (bottom). Right: a rotenone stimulus-specific eQTL for *ACSF3* in Sert (rs12597281, right). Shown are Manhattan plots for rotenone stimulated Sert at day 52 (top) and unstimulated day 52 Sert (bottom). (c) Comparison of the number of genes with at least one eQTL (number of eGenes; FDR < 5%; y-axis) as a function of effective sample size (number of unique donors; x-axis) across studies and cell types. Left: results from overlapping eQTL results in this study with *in vivo* eQTL maps from GTEx, divided into brain tissues and non-brain tissues. In red, the result from our study when aggregating across cell types and conditions. Right panel shows a magnified view of results from our study coloured by cell type and shaped by condition. (d) Sharing of eQTL signals discovered in our study for different cell types and conditions, as well as in two recent iPSC studies^2,34^ with *in vivo* brain eQTL maps (from GTEx). Violin plots show the extent of eQTL sharing **(Methods**), with each of 13 GTEx brain eQTL maps.

The largest number of eQTL were detected in progenitor cell populations, likely reflecting increased detection power due to the larger number of well-represented donors (>100 cells per donor; **Methods, Supplementary Fig. 8**). Notably, the cumulative number of genes with an eQTL in each cell type increased substantially when considering cells further progressed along the differentiation axis, as well as upon stimulation (**Fig. 4a**). For example, in DA cells, eQTL mapping in matured cells (day 52) identified an additional set of 270 egenes compared to day 30 cells.

An example of a timepoint-specific eGene is *HSPB1*, for which SNP rs6465098 is an eQTL in D52 cells, but not D30 (**Fig. 4b**). *HSPB1* encodes a heat shock protein that plays a key role in neuronal differentiation^36^ and for which changes in gene expression have been observed in neurons after ischemia37 and associated with toxic protein accumulation in Alzheimer disease^38,39^.

Similarly, we detected 196 additional eGenes with a rotenone-specific effect in DA and Sert neurons. As an example, rs12597281 is an eQTL for *ACSF3* in rotenone stimulated serotonergic neurons at day 52, but not in unstimulated cells (**Fig. 4b**). *ACSF3* encodes an acyl-CoA synthetase localized in the mitochondria and for which inherited mutations have been associated with a metabolic disorder, combined malonic and methylmalonic aciduria (CMAMMA), where patients exhibits a wide range of neurological symptoms including memory problems, psychiatric problems and/or cognitive decline^40^.

These examples highlight how changes in the expression of genes known to be associated with human disease can be transient and specific to a cell type and state. More importantly, this data shows how our experimental design brings an extra level of resolution to understand the disease mechanisms that were previously inaccessible from primary tissues and open up new experimental avenues.

To test how our eGene discovery relates to previous studies, we compared the number of eGenes identified in this study with bulk eQTL maps from *in vivo* tissues from the GTEx consortium^41^ (**Methods**). To aid the comparability between bulk and single-cell eQTL maps, we aggregated eQTL across cell types and found that the number of discovered eGenes was similar to that expected in a primary tissue of the same sample size (**Fig. 4c**). However, when focusing on individual cell populations, we observed fewer “cell type”-“condition” eQTL than detected in GTEx tissues of similar sample size, likely due to the decreased representation of donor cells.

A key question of eQTL maps from *in vitro* iPS-based models is how closely these resemble eQTL maps from primary tissues that differ in cell composition. To explore this, we tested the extent to which regulatory variants were shared between eQTL maps in three resources: 1) the current study, 2) GTEx brain tissues (n=13 tissues), and 3) bulk and single-cell RNA-seq profiles of HipSci iPS cell lines^2,34^, as measured by genome-wide consistency of eQTL effect sizes (using MASHR^42^; **Methods**). We found that as iPSCs were differentiated to increasingly mature neuronal cell types, the extent of eQTL sharing consistently increased (**Fig. 4d**). This provides additional confidence that eQTL discovered in iPS-derived neuronal populations mimic *in vivo* eQTL maps. Consistent with the trend of increased eQTL sharing, we also observed that the fraction of eQTL that are not represented in GTEx brain tissues decreases as the cells become more mature (**Supplementary Fig. 9a**). Interestingly, while iPS-derived eQTL maps mimic *in vivo* GTEx Brain eQTL maps, we also identified 2,203 eQTL that could not be detected in GTEx brain tissues (q-value>0.05 in any of 13 tissues, **Methods**), demonstrating the utility of iPSC and scRNA analysis to discover regulatory changes in disease associated genes.

### Co-localization of eQTL with disease risk variants

The identified cell-type specific eQTL maps across different differentiation contexts provide a unique opportunity to understand human disease traits and their genetic risk factors identified by genome-wide association studies (GWAS). To systematically test for such colocalization events, we applied COLOC^43^ (**Methods**) to the summary statistics from 25 neurological traits, eQTL discovered in our study, as well as eQTL obtained from GTEx (**Methods, Supplementary Table 8,9**).

In total, we identified 1,052 eQTL in our study with evidence of colocalization with at least one disease trait (**Fig. 5a,b**), 485 of which were found only in our data set. This corresponds to an additional ~10% of co-localization events of GWAS variants compared to eQTL across all GTEx tissues (5,028 across 48 tissues, **Fig. 5b**). Notably, 298 (61%) of the co-localizations in our data were associated with eQTL detected in later differentiation stages (D52) or upon stimulation (D52 ROT, **Supplementary Fig. 9b**).

**Figure 5.**
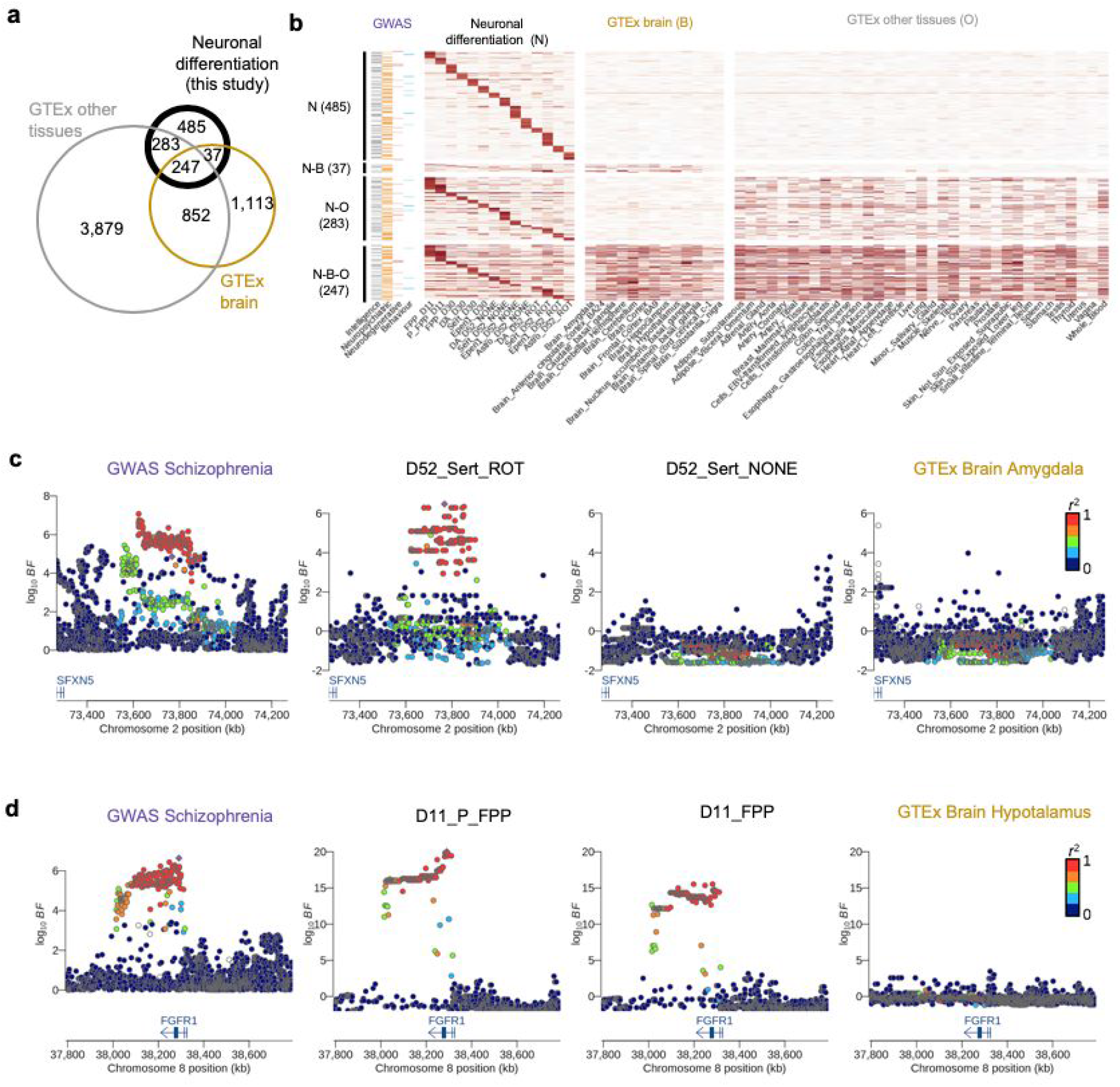
Colocalization analysis with 25 neuro-related GWAS traits. (a) Venn diagram showing the numbers of colocalization events overlapping between our study, GTEx brain and GTEx non brain tissues. (b) Heatmap showing the posterior probability of colocalization (PP4 from COLOC; Methods) for our eQTL that colocalised with one or more GWAS traits. N: Neuronal differentiation (this study), B: GTEx Brain, O: Other GTEx tissues. (c) Locus zoom plots around the *SFXN5* gene. The Schizophrenia GWAS association (left) is colocalised with the eQTL in rotenone stimulated serotonergic neurons at D52 (second panel from the left). No colocalization signal was detected in unstimulated serotonergic neurons at D52 (third panel from the left) or any other brain GTEx tissues as illustrated here with GTEx Brain Amygdala (rightmost panel). The lead variant is indicated with a purple diamond and other points were coloured according to the LD index (r2 value) with the lead variant. (d) A midbrain progenitor-specific eQTL for *FGR1* associated with schizophrenia. We identified a colocalization event with this eQTL in both proliferating (second panel from the left) and non-proliferating floor plate progenitors (third panel from the left) at day 11. No colocalization was found in any other cell type from our study (not shown) nor in any brain GTEx tissues (shown with GTEx Brain Hypothalamus, rightmost panel).

Among the most interesting colocalization events was an eQTL for *SFXN5*, a mitochondrial amino-acid transporter, which was specific to the rotenone-stimulated serotonergic neurons at day 52, and which co-localized with a Schizophrenia hit (PP4=0.76, **Fig. 5c**). Exposure to rotenone is known to induce oxidative stress by inhibiting the mitochondrial respiratory chain complex I^44,45^. We therefore speculate that the specific genetic signal observed for the mitochondrial gene *SFXN5* in serotonergic neurons is a possible factor modulating environmental stress response.

Another example that colocalized with a Schizophrenia GWAS variant was an eQTL for *FGFR1*, detected both in proliferating and non proliferating floor plate progenitors at D11 (PP4=0.93 and 0.88 respectively, **Fig 5d**). Previous studies have shown that nuclear *FGFR1* plays a key role in regulating neural stem cell proliferation and central nervous system development, in part, by binding to the promoters of genes that control the transition from proliferation to cell differentiation^46^. Additionally, it was shown that altered *FGFR1* signaling was linked to the progression of the cortical malformation observed in schizophrenia^47^.

These examples suggest that a combination of genetic and environmental factors during an early developmental stage might contribute to schizophrenia pathology and illustrate how these data represent a valuable resource to understand the molecular basis of complex neurological disease.

## Discussion

Characterising the function of human trait-associated genetic variation requires large scale studies performed in disease-relevant cell types and states. Here, we demonstrate how human iPSCs can be efficiently profiled at scale throughout a long-term differentiation to a midbrain cell fate. We uncover a highly reproducible, cell-intrinsic neuronal differentiation bias and show how this bias can be robustly predicted using simple gene expression profiling in the pluripotent cell state. This result sets the stage for the optimized design of future large scale iPSC experiments, where cell lines can be rationally selected *a priori* without the need for laborious testing of differentiation capacity. Importantly, our analysis also identified several hundred trait-associated genetic changes in gene expression that have not been previously identified. Overall, our study demonstrates the utility of pooled iPSC differentiation and single-cell analysis for revealing the function of disease-associated genetic variation in otherwise inaccessible cell states.

Despite a modest sample size, our study reveals a disproportionately large number of novel disease-eQTL colocalisations compared with GTEx tissues of equivalent sample size. For example, the number of novel disease-eQTL colocalisations added by GTEx liver or cerebellar hemisphere (n=208, 215 respectively) are 80 and 107, respectively, compared to 485 in this study. A simple explanation for this result is that our experiment profiled expression states that are hard to capture using post-mortem tissue, including timepoints during neuronal differentiation and rotenone exposure. Additionally, we detected many eQTL that were specific to individual cell types, enabled by the single-cell resolution of our study. These signals, while present, are challenging to detect in bulk tissue because the relevant cell types are often rare. Taken together, these results suggest that many “missing”, disease-relevant eQTL likely remain to be discovered using single cell sequencing of both primary tissue and *in vitro* cell models.

A second implication of our study is that, despite growth competition between cell lines, multiplexing experiments retain sufficient cells per donor to perform robust genetic analysis, even following extended periods in culture **(Supplementary Fig. 8, Supplementary Table 2**). Although cell lines were pooled at similar numbers, we observed extensive variation throughout our experiment in the numbers of cells produced by different lines. For example, 50% of the cells we sequenced were produced by only 12% of lines. Future technical improvements, such as more precise matching of growth rates of cell lines within pools, or line selection based on predicted differentiation capacity using markers in the iPS state may further increase the utility of multiplexed iPSC differentiation.

The “quality” of human iPSCs has previously been carefully examined using both genetic and functional genomic data^33,48–50^. Despite these efforts, differentiation bias among cell lines has been widely appreciated but poorly understood. The underlying mechanisms have been hypothesised to involve epigenetic factors, environmental factors such as culture conditions, or changes acquired by cells over time in culture, or cell type of origin. To the best of our knowledge, the work presented here is the first to systematically survey differentiation biases at the scale of an entire cell bank. To address this question, we leveraged the large number of cells in the HipSci bank and the detailed phenotyping of each of these lines. We excluded the cell type of origin hypothesis^51^ in this instance since all HipSci lines were skin-derived. We also observed relatively weak relationships between differentiation efficiency and other biological factors, such as X chromosome inactivation status, which has been described as relevant for other differentiation lineages^2^. Instead, we found that variability in differentiation outcomes can be largely explained by cell-intrinsic factors that are maintained over multiple freeze/thaw cycles. When we tested if these factors were due to the genetic variant inherited from the donor (^52^), we found that a strong donor component was unlikely due to the poor correspondence in predicted differentiation outcomes between lines derived from the same individual (**Supplementary Fig. 5c,d,e**). Additionally, we did not detect significant effects in a genome-wide association analysis with predicted differentiation outcomes (p>5×10^−8^, n=540, MAF=0.05). Given these results, we suggest the two most likely candidates for future investigation are somatic genetic changes or persistent changes to cell line epigenomes that occur early in cellular reprogramming.

In particular, our analysis indicates that the reduced production of neurons was best correlated with increased abundance of a specific subpopulation (cluster 2) of pluripotent cells that express the transcription factor *UTF1* and other genes at elevated levels. Counterintuitively, the proportion of cells in this subpopulation was positively correlated with the proportion of neuroblast cells on Day 11, but lower fractions of dopaminergic and serotonergic neurons at later stages of differentiation. One possible explanation is that cell lines that commit earlier to a neuronal fate disproportionately lose neurons upon passaging at Day 20. Alternatively, cluster 2 may preferentially give rise to radial glial cells that more readily switch to an astroglial and ependymal differentiation programme^53^. In support of this hypothesis, we identified several upregulated genes in cluster 2, including *SIX3, MT1F* and *PITX2*, that are thought to play a role in astrocyte and ependymal cell biogenesis^54–56^ (**Supplementary Table 4**). We speculate that culture methods that reduce iPSC heterogeneity may reduce the fraction of iPSC lines that resist efficient neuronal differentiation. We also note that our findings do not explain all of the variance in neuronal differentiation capacity, and future studies will be required to more fully understand the biological basis of the differentiation bias we have observed here.

Based on molecular markers that are predictive for differentiation bias, we estimate that 16% of iPSC lines in the HipSci resource produce few to no identifiable neuronal cell types under the conditions tested. While the production of neuronal cells was intrinsically limited in these cell lines, the fact that this effect was associated with particular cell lines but not with particular donors suggests that cell banks that contain multiple lines per donor can be most effectively utilised for applications involving neural differentiation by the rational selection of cell clones. Importantly, this *a priori* selection is enabled by gene expression profiling data from the pluripotent state that is easily obtainable and often already available.

In summary, our study clearly demonstrates how iPSC differentiation combined with single cell RNA-seq unlocks population level studies in increasingly complex, dynamic and biologically realistic cellular models. We anticipate that future uses of this model system will focus on experimental settings that are challenging or impossible with primary cells. These could include high resolution sampling along extended differentiation times to more complex differentiation trajectories, such as cell organoids, or involve large panels of disease relevant-stimuli and drug exposures. Collectively, our study will guide future efforts to understand the common genetic basis of neurological disorders, and facilitate the development of iPSC-based approaches for modelling and treating these diseases.

## Methods

### Human iPSC culture

Feeder-free human iPSCs were obtained from the HipSci project^1^. Lines were thawed onto tissue culture-treated plates (Corning, 3516) coated with 10 μg/mL VitronectinXF (StemCell Technologies, 07180) using complete Essential 8 (E8) medium (Thermo Fisher, A1517001) and 10 μM Rock inhibitor (Sigma, Y0503). Cells were expanded in E8 medium for 2 passages using 0.5 μM EDTA pH 8.0 (Thermo Fisher, 15575-020) for cell dissociation.

### Pooling and differentiation of midbrain dopaminergic neurons

iPSC colonies were dissociated into a single-cell suspension using Accutase (Thermo Fisher, A11105-01) and resuspended in E8 medium containing 10 μM Rock inhibitor. Cells were counted using an automated cell counter (Chemometec NC-200) and a cell suspension containing an equal amount of each iPSC line was prepared in E8 medium containing 10 μM Rock inhibitor and seeded at 2 x 10^5^ cells per cm^2^ on 0.01% Geltrex-(Thermo Fisher, A1413202) coated plates. Each pool of lines contained between 7 to 24 donors. 24h after plating, neuronal differentiation of the pooled lines to a midbrain lineage was performed as described by^12^ with minor modifications: 1. SHH C25II was replaced by 100nM SAG (Tocris, 6390) in the neuronal induction phase. 2. On day 20, the cells were passaged with Accutase containing 20 units/mL of papain (Worthington, LK00031765) and plated at 3.5 x 10^5^ cells per cm^2^ on 0.01% Geltrex-coated plates for final maturation.

### Rotenone stimulation

On day 51 of differentiation, the cells were exposed for 24h to freshly prepared 0.1 μM rotenone (Sigma, R8875, purity HPLC ≥ 95%) diluted in neuronal maturation medium^12^. The final DMSO concentration was 0.01% in all exposure conditions. Unstimulated control samples (i.e. DMSO only) were taken concurrently.

### Generation of cerebral organoids

Cerebral organoids were generated according to the enCOR method as previously described by^32^. Briefly, one pool of 18 iPSC lines was thawed and expanded for 1 passage before seeding 18,000 cells onto PLGA microfilaments prepared from Vicryl sutures. STEMdiff Cerebral Organoid kit (Stem Cell Technologies, 08570) was used for organoid culture with timing according to manufacturer’s suggestion and Matrigel embedding as previously described^57^. From day 35 onward the medium was supplemented with 2% dissolved Matrigel basement membrane (Corning, 354234), and processed for scRNA-seq after 113 days of culture.

### Generation of single cell suspensions for sequencing

On harvesting days, the cells were washed once with 1X DPBS (Thermo Fisher, 14190-144) before adding either Accutase (day 11) or Accutase containing 20 units/mL of papain (days 30 and 52). The cells were incubated at 37°C for up to 20 min (day 11) or up to 35 min (days 30 and 52) before adding DMEM:F12 (Thermo Fisher Scientific, 10565-018) supplemented with 10 μM Rock inhibitor and 33 μg/mL DNase I (Worthington, LK003170, only for days 30 and 52). The cells were dissociated using a P1000 and collected in a 15 mL tube capped with a 40 μm cell strainer. After centrifugation, the cells were resuspended in 1X DPBS containing 0.04% BSA (Sigma, A0281) and washed 3 additional times in 1X DPBS containing 0.04% BSA. Single-cell suspensions were counted using an automated cell counter (Chemometec NC-200) and concentrations adjusted to 5 x 10^5^ cells/mL.

Organoids were washed twice in 1X DPBS before adding EBSS (Worthington, LK003188) dissociation buffer containing 19 U/mL of papain, 50 μg/mL of DNase I and 22.5X of Accutase. Organoids were incubated in a shaking block (750 rpm) at 37°C for 30 min. Every 10 min, the organoids were triturated using a P1000 and BSA-coated pipette tips until large clumps were dissociated. Dissociated organoids were transferred into a new tube capped with a 40 μm cell strainer and pelleted for 4 min at 300g. After centrifugation, the cells were resuspended in EBSS containing 50μg/mL of DNase I and 2 mg/mL ovomucoid (Worthington, LK003150). 0.5 volume of EBSS, followed by 0.5 volume of 20 mg/mL ovomucoid were added to the top of the cell suspension and the cells were mixed by flicking the tube. After centrifugation, the cells were resuspended in 1X DPBS containing 0.04% BSA. Single-cell suspensions were counted using an automated cell counter and concentrations adjusted to 5 x10^5^ cells/mL.

### Immunohistochemistry

Cells were fixed in 4% paraformaldehyde (Thermo Fisher Scientific, 28908) for 15 min, rinsed 3 times with PBS1X (Sigma, D8662) and blocked with 5% normal donkey serum (NDS; AbD Serotec, C06SBZ) in PBST (PBS1X + 0.1% Triton X-100, Sigma, 93420) for 2h at room temperature. Primary antibodies were diluted in PBST containing 1% NDS and incubated overnight at 4°C. Cells were washed 5 times with PBS1X and incubated with secondary antibodies diluted in PBS1X for 45 min at room temperature. Cells were washed 3 more times with PBS1X and Hoechst (Thermo Fisher Scientific, H3569) was used to visualize cell nuclei. Image acquisition was performed using Cellomics array scan VTI (Thermo Fisher Scientific).

The following antibodies were used:

FOXA2 (Santa Cruz, sc101060 - 1/100)

LMX1A (Millipore, AB10533 - 1/500)

TH (Santa Cruz, sc-25269 - 1/200)

MAP2 (Abcam, 5392 - 1/2000)

Donkey anti-chicken AF647 (Thermo Fisher Scientific, A21449)

Donkey anti-mouse AF488 (Thermo Fisher Scientific, A11008)

Donkey anti-mouse AF555 (Thermo Fisher Scientific, A31570)

Donkey anti-rabbit AF488 (Thermo Fisher Scientific, A21206)

Donkey anti-rabbit AF555 (Thermo Fisher Scientific, A27039)

### Chromium 10x Genomics library and sequencing

Single cell suspensions were processed by the Chromium Controller (10x Genomics) using Chromium Single Cell 3’ Reagent Kit v2 (PN-120237). On average, 15,000 cells from each 10x reaction were directly loaded into one inlet of the 10x Genomics chip (**Supplementary Table 1**). All the steps were performed according to the manufacturer’s specifications. Barcoded libraries were sequenced using HiSeq4000 (Illumina, one lane per 10x chip position) with 50bp or 75bp paired end reads to an average depth of 40,000-60,000 reads per cell.

### Single-cell data pre-processing

Sequencing data generated from the Chromium 10x Genomics libraries (see above) were processed using the CellRanger software (version 2.1.0) and aligned to the GRCh37/hg19 reference genome. Counts were quantified using the CellRanger “count” command, with the Ensembl 84 reference transcriptome (32,738 genes) with default QC parameters.

For each of 19 pooled experiments, donors (i.e. cell lines) were demultiplexed using demuxlet^15^, using genotypes of common (MAF>1%) exonic variants available from the HipSci bank, and a doublet prior of 0.05. Only single cells with successful donor identification were retained for further analysis. This step filtered out two types of droplet: those containing two or more cells from different individuals, and those containing no cells, but that contained a mixture of free-floating RNA and had therefore passed the CellRanger UMI filter (above).

Further quality control was applied to exclude eight 10x samples, where a 10x sample is defined as the cells sequenced from one inlet of a 10x chip. In particular: samples for pool 10 on Day 11 were excluded because of an issue in the library preparation. Samples for pool 12 on Day 52 were excluded on the basis of low cell viability (72.1% in the rotenone-stimulated sample), and outlying gene expression (with the first principal component in gene expression separating this sample from others at the same time point). The two 10x reactions for pool 13 on Day 30 were excluded on the basis of low quality metrics, likely as a result of cell overloading (~25,000 cells loaded). Finally, cells from an outlier cell line (HPSI0913i-gedo_3) were excluded. This cell line contributed a very high proportion of cells to samples from pool 14 (91%), and had outlying gene expression, raising concerns that this cell line had acquired a large-effect somatic mutation (**Supplementary Table 1**).

### Normalisation, dimensionality reduction, and clustering

Two sets of analyses were performed: i) analysis of each time point independently, ii) a combined analysis of a subsample of 20% of cells from all time points (used only for visualisation purposes; **Fig. 1**).

First, independent analysis of time points allowed efficient batch effect correction (as all samples were from the same time point, containing similar mixtures of cell types), as well as reducing computational demands (by reducing the number of cells analysed together). For the analysis of each time point independently, the following steps were performed: counts were normalised to the total number of counts per cell. Only genes with non-zero counts in at least 0.5% of cells were retained. The top 3,000 most variable genes were then selected, after controlling for mean-variance dependence in expression data. The first 50 principal components (PCs) were calculated. Batch correction was applied on the level of PCs using Harmony^16^, with each 10x sample treated as a distinct batch. UMAP and clustering was performed using these transformed PCs. Clustering was performed using Louvain clustering with 10 nearest neighbours. Analysis steps besides batch correction were carried out using the Scanpy package^58^. Clusters were mapped to cell types using a set of literature-curated 48 marker genes of major brain cell types. When two clusters show the same gene set enrichment they were assigned the same cell type identity (**Supplementary Fig. 2**).

Second, for the combined analysis of all time points (for visualization purposes only), the same steps were applied, except that only a random subsample of 20% of cells were included in the analysis (following filtering for cells with donor assignment), and the definition of batches for the Harmony batch correction step. In particular, in order for each batch to have a similar mixture of cell types, each pool (rather than each 10x sample) was considered as a distinct batch.

### Batch correction and clustering of the organoid dataset

The same steps of dimensionality reduction, batch correction and clustering described above were applied to the cerebral organoid data. This identified eight clusters that were mapped to different cell types (neural cells, intermediate progenitor cells, radial glial progenitor cells, satellite cells, mesenchymal cells, myotube and Wnt and PAX7 positive cells) using 24 marker genes (**Supplementary Fig. 4**).

### Batch correction and clustering of the two single cell iPSCs datasets

The same dimensionality reduction, batch correction and clustering steps described above were applied to the two single cell iPSC datasets analysed^2,35^ (**Supplementary Fig. 6,7**). For the Cuomo et al dataset^2^, normalised (by CPM) and log-scaled data were taken from the original publication and no further normalisation was performed. For the Sarkar et al dataset^35^, count data were normalised and log-scaled as described above for our data (normalised to total counts per cell). Note that in both cases only QC-passing cells (as defined in the original publications) from these datasets were included. This analysis identified five and four clusters in the two different datasets, respectively.

### Definition of differentiation efficiency

We computed cell type proportions for each cell line in each pool (i.e. all (cell line, pool) combinations) at each time point. Based on these proportions, (cell line, pool) combinations were clustered, based on Euclidean distance (**Fig. 3b, Supplementary Fig. 3**). Only (cell line, pool) combinations for which at least 10 cells were present at all time points were included in the heatmap in **Supplementary Fig. 3a** and in the PCA analysis shown in **Supplementary Fig. 3**.

Differentiation efficiency was defined as the sum of the proportion of serotonergic and dopaminergic neurons present on Day 52. The differentiation efficiency of iPSC lines was calculated as the average of the efficiencies across all pools in which that cell line was included. That is, for those iPSC lines which were included in multiple pools, differentiation efficiency was averaged across the results from differentiation in independent pools (replicates, as in **Fig. 2b**).

### Predictive model of differentiation efficiency from iPSC gene expression

A logistic regression classifier was trained to predict midbrain neuron differentiation failure (differentiation efficiency < 0.2, from above definition) of iPSC lines from their gene expression at the iPSC stage. For cell lines that were differentiated multiple times, their differentiation efficiency was taken as the mean of its differentiation efficiencies in the replicate experiments. Gene expression data was available from independent bulk RNA-seq experiments^34^. The feature set was all expressed genes (i.e. genes with mean log2(TPM+1)>2, n=13,475). The model was trained and tested using the scikit-learn (v0.21.3) package in Python, with the function sklearn.linear_model.LogisticRegression. L1 regularisation was used, with default parameter settings (inverse regulation strength = 1.0). Precision-recall was evaluated using leave-one-out cross validation. That is, for each data point in turn, a model was trained on all other data, and its ability to predict the left out data point was evaluated.

When predicted scores for two lines from the same donor were present, we classified donors into concordant good differentiators when both lines scored > 0.5 (n=183), concordant bad differentiators when both lines scored < 0.5 (n=25). Finally, “discordant” donors were donors for which the two lines scored differently, one >0.5, one <0.5 (n=62). Bulk RNA-seq expression of *UTF1* and *TAC3* for these lines confirmed such predictions (**Supplementary Fig. 5d**)

### X chromosome inactivation (XCI) status

XCI was assessed by considering allele-specific expression (ASE) from the X chromosome, as quantified by bulk RNA-seq. Allele-specific counts were obtained for SNPs present in DBSNP using GATK ReadCounter with the command ‘GenomeAnalysisTK.jar -T ASEReadCounter -U ALLOW_N_CIGAR_READS --minMappingQuality 10 --minBaseQuality 2’. Heterozygous SNPs located on the X chromosome for which the total number of overlapping reads was > 20 were retained for analysis. For each SNP, the ASE fraction was defined as the fraction of reads mapping to the less expressed allele (thus the ASE fraction was ≤ 0.5 for all SNPs). For each sample, the XCI status was quantified as the mean ASE fraction of all heterozygous X chromosome SNPs in that sample.

### Association of cell line metadata features with differentiation efficiency

We tested for associations between differentiation efficiency and the cell line donor’s sex (n=199, t-test) as well as XCI status (on the subset of female lines; n=97), passage number (n=195), two pluripotency scores (n=196, F-test). The pluripotency scores used were the pluritest score and the novelty score, which were generated in the course of banking the HipSci cell lines^33^.

Next, we tested the same features for associations with the predicted differentiation scores we estimated for a larger set of HipSci lines as described above (**Supplementary Table 5**). Again, we tested for associations with a cell line’s donor sex (n=812, t-test) as well as XCI status (n=342), two pluripotency scores (n=797, F-test). We note that, unlike in our experiments, this dataset included cell lines in both feeder and feeder-free culture conditions, and therefore we tested this as an additional factor denoted ‘feeder free status’ (n=812, t-test).

We corrected for multiple testing separately for the two sets of tests performed using Bonferroni correction. Results from this analysis are in **Supplementary Table 3**.

### Differential expression analysis

Differentiation expression analysis between each cluster and the others in the single cell iPSC datasets was performed using scanpy’s function “tl.rank_genes_groups”, grouping by each cluster at a time^58^. Briefly, the function computes Z-scores as well as log fold changes. Nominal p-values are computed using a t-test like test, using approximated log mean values, whilst adjusted p-values using Benjamini-Hochberg. We report significant results at FDR < 5% (**Supplementary Table 6**).

### *cis* eQTL mapping

For *cis* eQTL mapping, we followed Cuomo et al^2^, and adopted a strategy similar to approaches commonly applied in conventional bulk eQTL analyses^1^. We considered common variants (minor allele frequency (MAF) > 5%) within a *cis-* region spanning 250kb up- and downstream of the gene body for *cis* QTL analysis. Association tests were performed adapting a linear mixed model (LMM), previously used for single cell *cis* eQTL mapping^2^. However, instead of accounting for population structure we used a random effect term to account for varying numbers of cells per donor. Briefly, for each donor we introduced a variance term 1/n, accounting for the varying numbers of cell used to estimate mean expression level for each donor. All models were fitted using LIMIX^59,60^. The values of all features were standardized and the significance was tested using a likelihood ratio test (LRT). To adjust for experimental batch effects across samples, we included the first 15 principal components calculated on the expression values in the model as fixed effect covariates. These batch effects usually affect the expression of many genes, and therefore are detectable in the principal components of expression. Furthermore, such global batch effects are orthogonal to the effects of a single cis regulatory variant on the expression of one gene. In order to adjust for multiple testing, we used an approximate permutation scheme, analogous to the approach proposed in^61^. Briefly, for each gene, we generated 1,000 permutations of the genotypes while keeping covariates, random effect terms, and expression values fixed. We then adjusted for multiple testing using this empirical null distribution. To control for multiple testing across genes, we then applied the Storey’s Q value procedure^61,62^). Genes with significant eQTL were reported at an FDR < 5%.

We performed eQTL mapping as described above for each of the well represented (top 4 cell type per condition with at least 20% cells) contexts (cell type, time point and stimulus status, **Supplementary Table 6**). Gene expression for each donor was calculated as the mean of log-transformed counts-per-cell-normalised expression across cells (including cells from different pools, where applicable). Each individual quantification (i.e. expression of a donor in a particular cell type) was only included in the eQTL mapping analysis if it represented the mean of >10 cells. For each context (cell type, condition), all genes detected in at least 1% of cells of that context were included (this varied from 11,659 to 13,536 genes tested). The minimum P-value SNP per gene (lead eQTL variant) is reported in **Supplementary Table 6**.

We also performed cis eQTL mapping for all 14 contexts without using the noise matrix accounting for the number of cells, resulting in a substantially smaller number of discoveries (**Supplementary Figure 8b**).

### Sharing of eQTL signal with GTEx brain tissues

To quantify the amount of sharing between each two pairs of eQTL maps (our cell type-condition maps to each of 13 eQTL maps of brain tissues from GTEx) we used the MASHR software^42^. Briefly, the effect sizes (betas) were calculated for each SNP-gene pair across the 14 cell type-condition eQTL maps from our study and extracted from the 13 brain tissues from the GTEx catalogue, as well as two iPSC eQTL maps, using scRNA-seq and bulk RNA-seq respectively^2,34^. Only genes expressed in all GTEx tissues, in iPSC^34^ and all of our contexts were included (n=6,205). To calculate the sharing of effects we selected the top eQTL SNP per gene, based on the effect sizes in the tissues with more than 150 samples. Four random SNPs per gene were selected as a background for the calculation of the data driven covariance; we also included a canonical covariance matrix as recommended^63^. Next, we extracted the posterior betas, using MASHR, and estimated pairwise levels of sharing between conditions/tissues where we defined shared as “the same sign and effect size within a factor 0.5 of each other”.

### Colocalization analysis between neuro-related GWAS traits

We collected summary statistics for 25 GWAS traits that were either neurodegenerative/ neuropsychiatric diseases or related to behaviour and intelligence, and that have at least 5 GWAS loci with the significance threshold of p=5.0×10^−8^ (**Supplementary Table 8**). We then defined GWAS subthreshold loci as 1Mb-wide genomic windows with at least one SNP at *P*-value<10e-6, centring the window around the index variant (variant with minimum *P*-value in the window). If there were multiple subthreshold loci within a 1Mb window, we merged them and took the index variant with the minimum *P*-value overall. Statistical colocalization analysis between 14 eQTL maps from our study and 48 eQTL maps from GTEx (v7) and those GWAS loci was performed using the COLOC package^43^, implemented in R with default hyperparameter setting. We tested any gene whose transcription start site (TSS) and eQTL lead variant (minimum P-value SNP for the gene) were both within the 1Mb window centered at each GWAS index variant. We tested all SNPs located between the GWAS index variant and the top eQTL (within the window) variant with 500Kb extensions either side. We matched SNPs between eQTL and GWAS based on chromosomal position and reference/alternative alleles. Genes with the posterior probability of colocalization (PP4) greater than 0.5 were defined as GWAS colocalization.

## Data availability

Single cell RNA-seq data will be made available soon.

## Code availability

All scripts are available in the following github repository: https://github.com/single-cell-genetics/singlecell_neuroseq_paper/

The eQTL mapping pipeline is available here: https://github.com/single-cell-genetics/limix_qtl/

## URLs

HipSci: http://www.hipsci.org. GTEx: https://www.gtexportal.org/home/datasets

## Acknowledgements

All data for this study was generated under Open targets project OTAR039. J.J was supported by a postdoctoral fellowship from OpenTargets, ASE.C was supported by a PhD fellowship from the EMBL International PhD Programme (EIPP) and DD.S was supported by a postdoctoral fellowship from EMBL Interdisciplinary Postdoc (EIPOD) programme. N.K and DJ.G. were funded by the Wellcome Trust grant WT206194. FTM is a New York Stem Cell Foundation - Robertson Investigator and is supported by The New York Stem Cell Foundation [NYSCF-R-156], the Wellcome Trust and Royal Society [211221/Z/18/Z], and the Chan Zuckerberg Initiative [191942]. JC.M acknowledges core support from EMBL and Cancer Research UK. Research in the Stegle lab is supported by the BMBF, the Volkswagen Foundation and the EU (ERC project DECODE). We thank the MRC Metabolic Diseases Unit Imaging Core Facility for assistance with imaging. We thank the staff in the Cellular Genetics and Phenotyping and Sequencing core facilities at the Wellcome Sanger Institute. We thank Helena Kilpinen and Pau Puigdevall Costa for the very useful discussions regarding data analysis.

## Author contributions

The main analyses and data preparations were performed by JJ, DD.S and ASE.C; NK performed the colocalization analysis. Cell culture experiments were performed by JJ, JH, JS, DP and MA and MA.L performed the experimental work on the organoid dataset. JJ, DD.S, ASE.C, JM, FT.M, OS and DJ.G wrote the manuscript; N.K assisted in editing the manuscript; J.J, FT.M, OS and DJ.G conceived and oversaw the study.

