## Supplementary Information for "Population-scale single-cell RNA-seq profiling across dopaminergic neuron differentiation"

Jerber *et al.*

**Overview of Supplementary Figures:**

Supplementary Figure 1: Immunostaining of midbrain neural progenitors and dopaminergic neurons.

Supplementary Figure 2: Clustering and cell type assignment.

Supplementary Figure 3: Cell proportion distribution at day 52 leads to defining differentiation efficiency.

Supplementary Figure 4: Cerebral organoid analysis.

Supplementary Figure 5: Predicting differentiation failure from iPSC gene expression.

Supplementary Figure 6: Re-analysis of iPSC scRNA-seq data reveals a subpopulation characterised by expression of predictive marker genes associated with lower differentiation efficiency.

Supplementary Figure 7: Analysis of a single cell iPSC dataset from Sarkar *et al.*

Supplementary Figure 8: Number of cells per cell type and relation to eQTL power.

Supplementary Figure 9: eQTL and colocalisation (extended).

**Overview of Supplementary Tables:**

Supplementary Table 1: Overview of the collected samples

Supplementary Table 2: Cell distribution per cell line, pool, cell type, time point and stimulation

Supplementary Table 3: Associations between cell lines' factors and differentiation efficiency

Supplementary Table 4: Liof genes correlated with differentiation efficiency

Supplementary Table 5: Predictive scores for HipSci banked lines

Supplementary Table 6: DE genes between cluster 2 and others (single cell iPSC data).

Supplementary Table 7: List of eQTL discoveries at FDR 5%

Supplementary Table 8: List of neurological traits used for colocalisation analysis

Supplementary Table 9: Results of the colocalisation analysis

**Supplementary Figure 1.** Immunostaining of midbrain neural progenitors and dopaminergic neurons. (a) Immunostaining for known midbrain progenitor markers LMX1A and FOXA2 at day 11. Nuclei were counterstained with Hoechst. Scale bar: 25µm. (b) Immunostaining of differentiated dopaminergic neurons for the neuronal marker MAPT2 (white) and the dopaminergic neuronal markers TH and LMX1A. Scale bar: 25µm. Data is shown for two example individual cell lines (HPSI0155i-hecn_6 and HPSI0514i-uenn_3) as well as three entire differentiation pools (Pools 1,2,3).

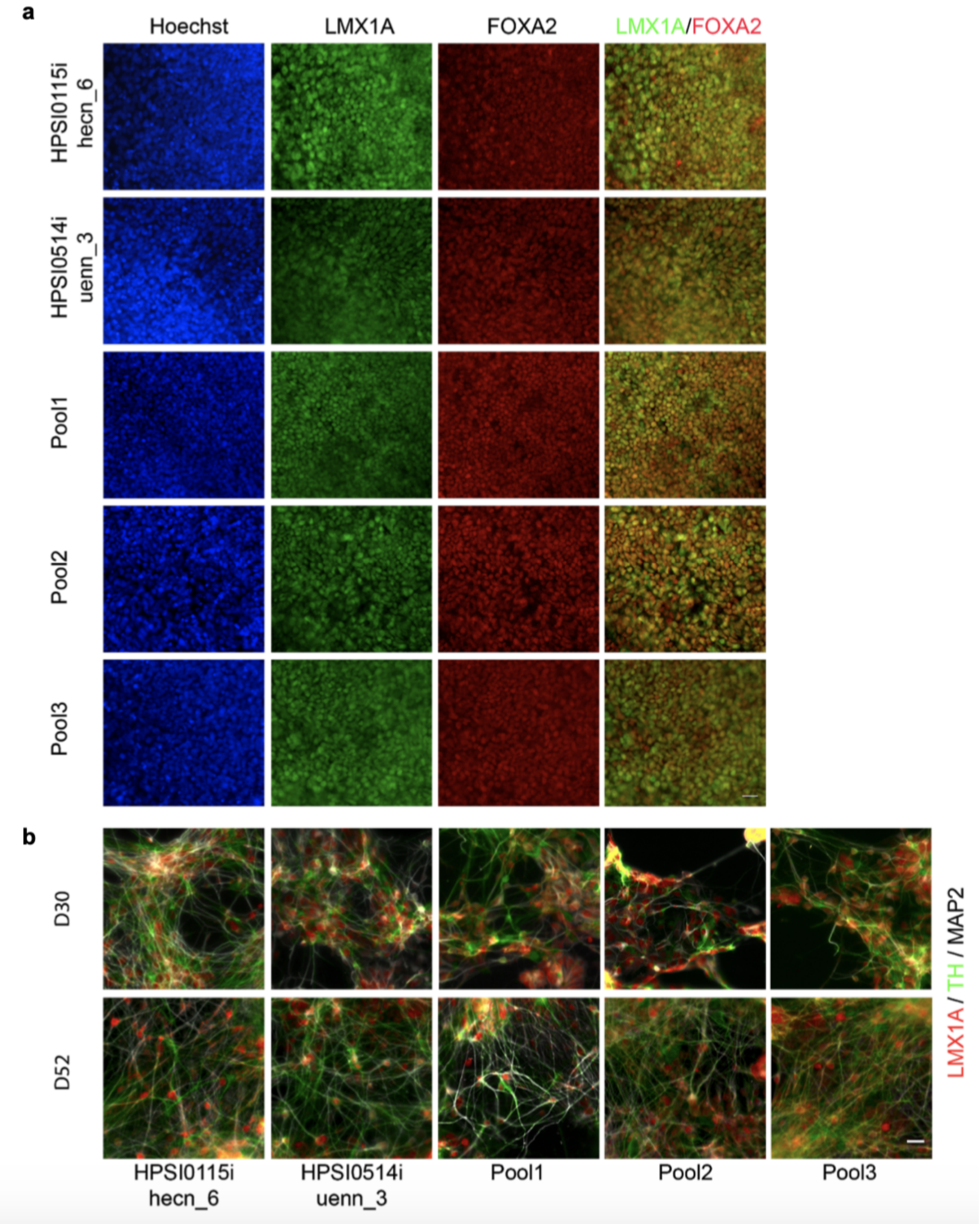

**Supplementary Figure 2.** Clustering and cell type assignment. Cells at each time point were clustered using Louvain clustering, after normalization and batch correction using Harmony[1]. Subsequently, clusters were assigned to cell types using known marker genes. When two clusters showed the same gene set enrichment they were computationally assigned to the same cell type identity (Methods). (a) UMAPs of cells sampled at each time point and coloured by cell clusters. (b) UMAPs of cells sampled at each time point and coloured by assigned cell types. (c) Heatmap showing the expression profile of canonical neuronal marker genes across the identified cell types. Astro: Astrocytes-like, DA: Midbrain dopaminergic neurons, Epen1/2: Ependymal-like 1/2, FPP: Floor Plate Progenitors, NB: Neuroblasts, P_FPP: Proliferating Floor Plate Progenitors, P_Sert: Proliferating serotonergic neurons, Sert: Serotonergic neurons, U_Neur: Unknown Neurons.

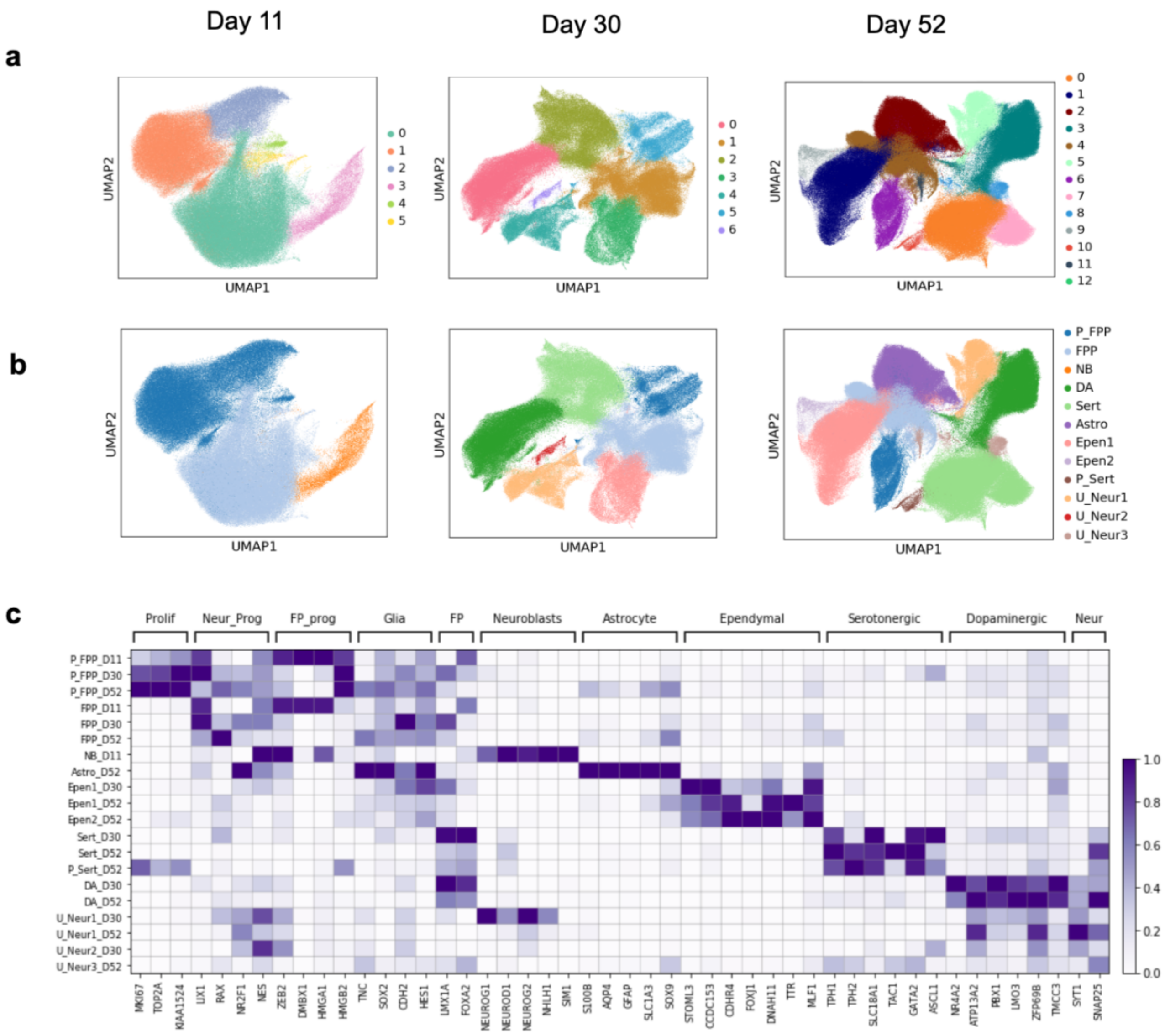

**Supplementary Figure 3.** Distribution of cell proportion at day 52 and the definition of differentiation efficiency. Cell proportions were generated for each cell type and time point for all combinations of cell lines and pools with at least 10 cells at all time points (10 pools). (a) Heatmap of the resulting cell proportion matrix. Pools are shown in the first bar and the colours indicate in which of the 10 pools each line was differentiated. Rows (i.e. cell line, pool combinations) were hierarchically clustered according to Euclidean distance (as in **Fig. 2b**). (b) Proportion of variance explained by each principal component calculated from cell proportions matrix. (c) Comparison of the first principal component (PC1) to the sum of fractions of dopaminergic and serotonergic neurons present on Day 52. (d) UMAP of the corresponding expression dataset (considering cells as in a), with cells coloured according to the principal components from the cell proportion matrix described above.

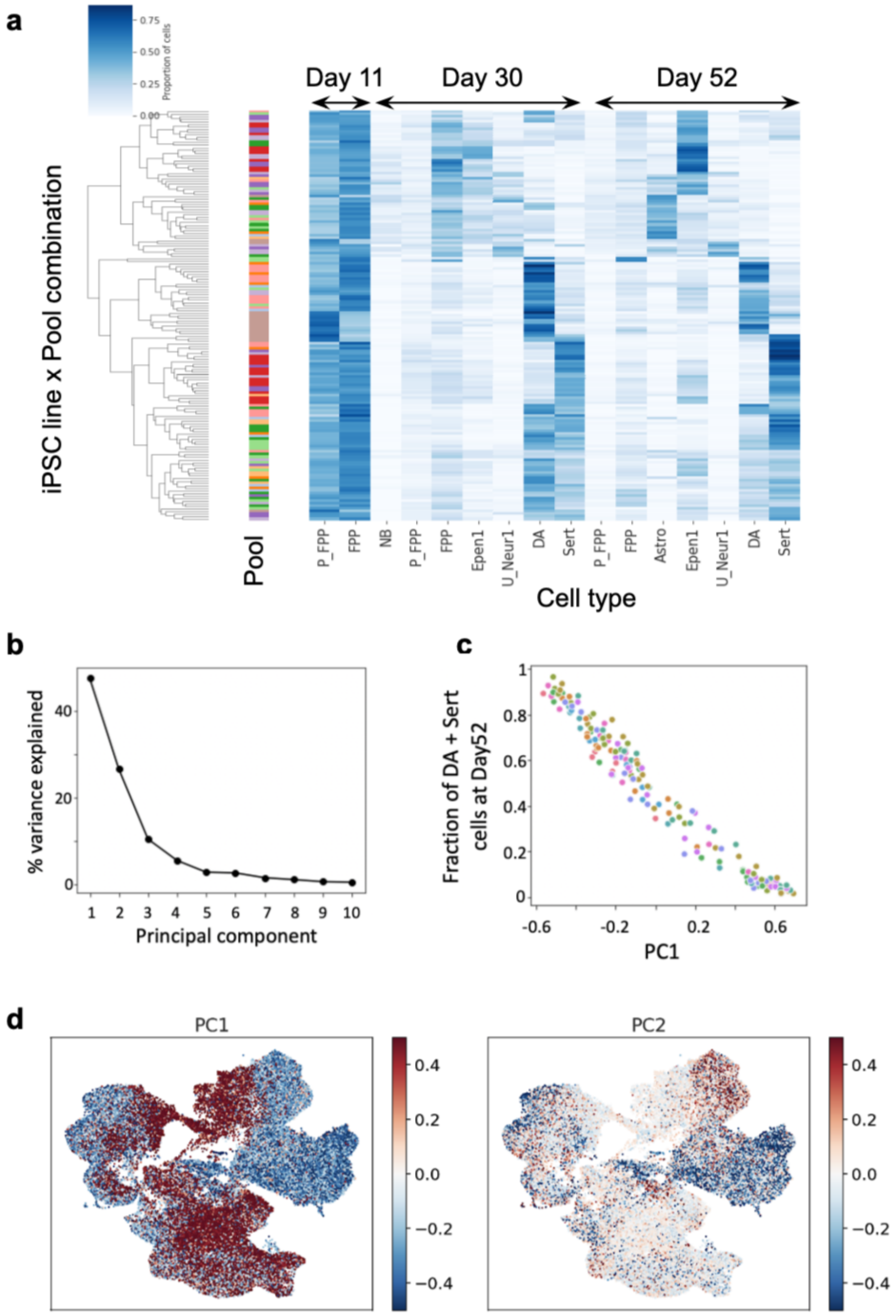

**Supplementary figure 4.** Cerebral organoid analysis. (a) UMAP of cerebral organoid cells, as assayed by scRNA-seq after 113 days of differentiation, coloured by cell lines. (b) Matrix plot showing the expression profile of genes known to be expressed in each of the identified cell types. IP: Intermediate progenitors, RGP: Intermediate glial progenitors, PAX7+ Satellite: Satellite cells PAX7+, Mesenchymal: Mesenchymal cells, Wnt+: Wnt_positive cells.

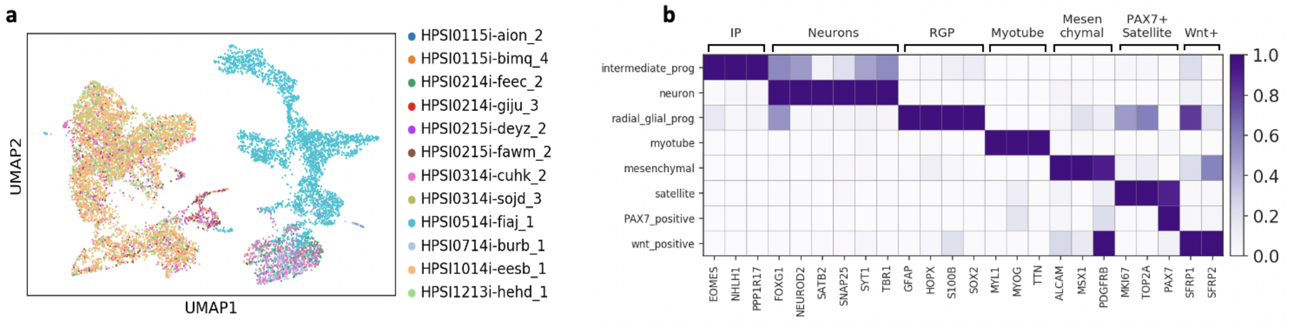

**Supplementary Figure 5.** Predicting differentiation failure from iPSC gene expression. (a) Histogram of differentiation efficiencies across cell lines. The threshold chosen to define differentiation success or failure (i.e. efficiency=0.2) is shown by the dashed line, separating the two modes of the distribution. (b) Precision-recall curves for a logistic regression model predicting differentiation failure from iPSC gene expression data[2] using a range of thresholds between 0.1 and 0.35 to define differentiation failure. Results are presented from leave-one-out cross validation (Methods). (c) Histogram of predicted differentiation scores across all HipSci cell lines. The bimodal distribution is especially extreme in this case, and 0.5 was used as the threshold to split bad from good differentiator lines. (d) Scatterplot of predicted differentiation scores for donors for which we have data for two different cell lines. Replicate1 (rep1) is chosen as the line with lower predicted score (n=270). Colours indicate three categories of donors, according to whether both lines from the same donor are predicted to fail differentiation (blue), both are predicted to succeed (green), the two lines are discordant (one is predicted to successfully differentiate, but not the other, yellow). (e) Bulk RNA-seq expression of *UTF1, TAC3* for the two replicate lines per donor stratified by the categorization described in (d).

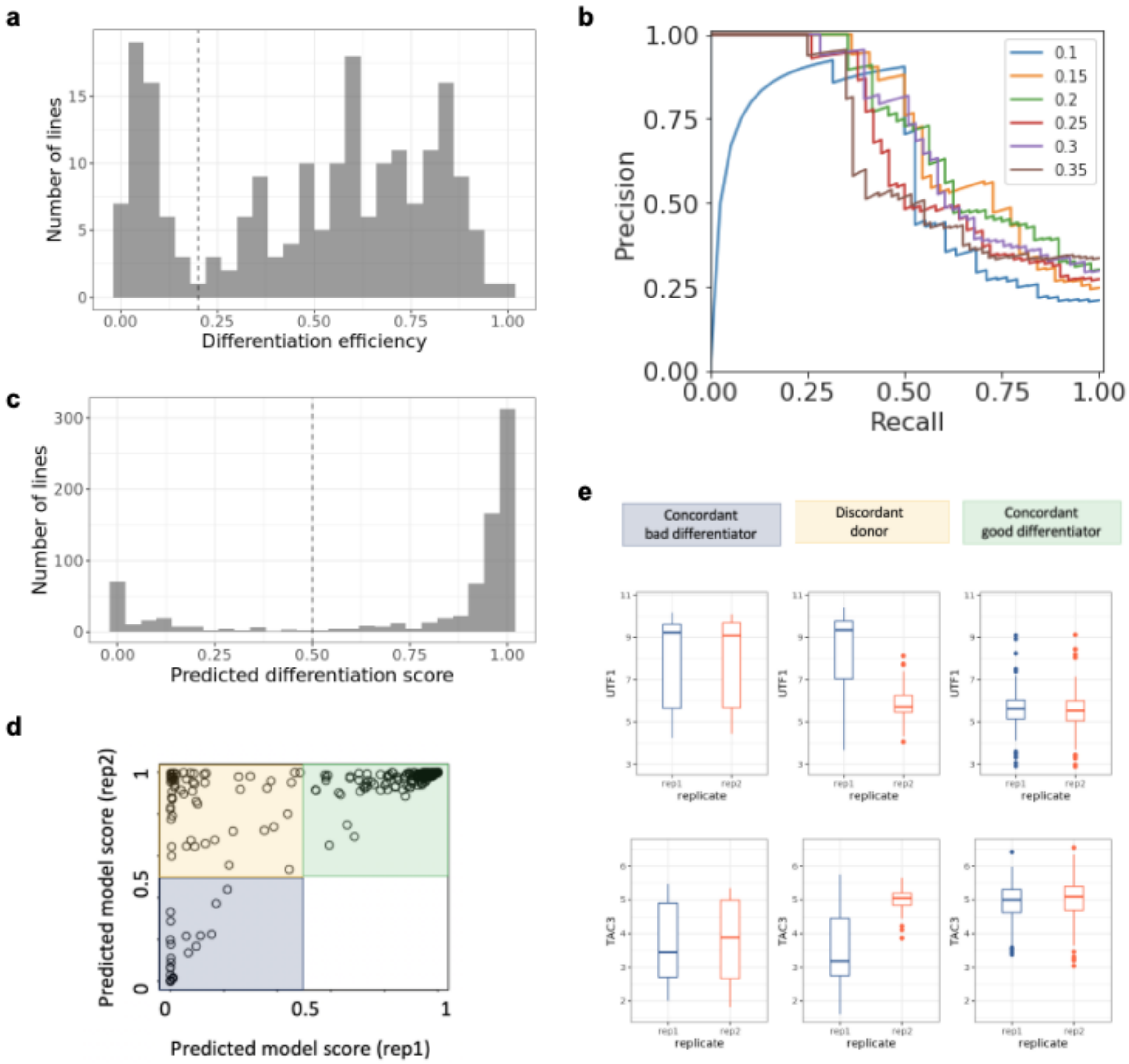

**Supplementary Figure 6.** Re-analysis of iPSC scRNA-seq data reveals a subpopulation characterised by expression of predictive marker genes associated with lower differentiation efficiency. (a) UMAP overview of the dataset. iPSC scRNA-seq data from [3] were re-analysed following the same batch correction and clustering steps applied to our neural differentiation data, identifying 5 clusters. (b) Violin plots of gene expression for genes related to pluripotency (*NANOG, SOX2, POU5F1*) and two gene markers that are respectively upregulated and downregulated in cluster 2 (*UTF1, TAC3,* from **Fig. 3**). (c) Scatter plot showing the proportion of cells assigned to cluster 2 between replicates (n=23). (d) Scatterplot between the proportion of cells assigned to cluster 2 (y-axis) and differentiation efficiency (x-axis) similar to **Fig 3.f** but where we use imputed proportions of cluster 2 cells from bulk RNA-seq available for most cell lines (n=182).

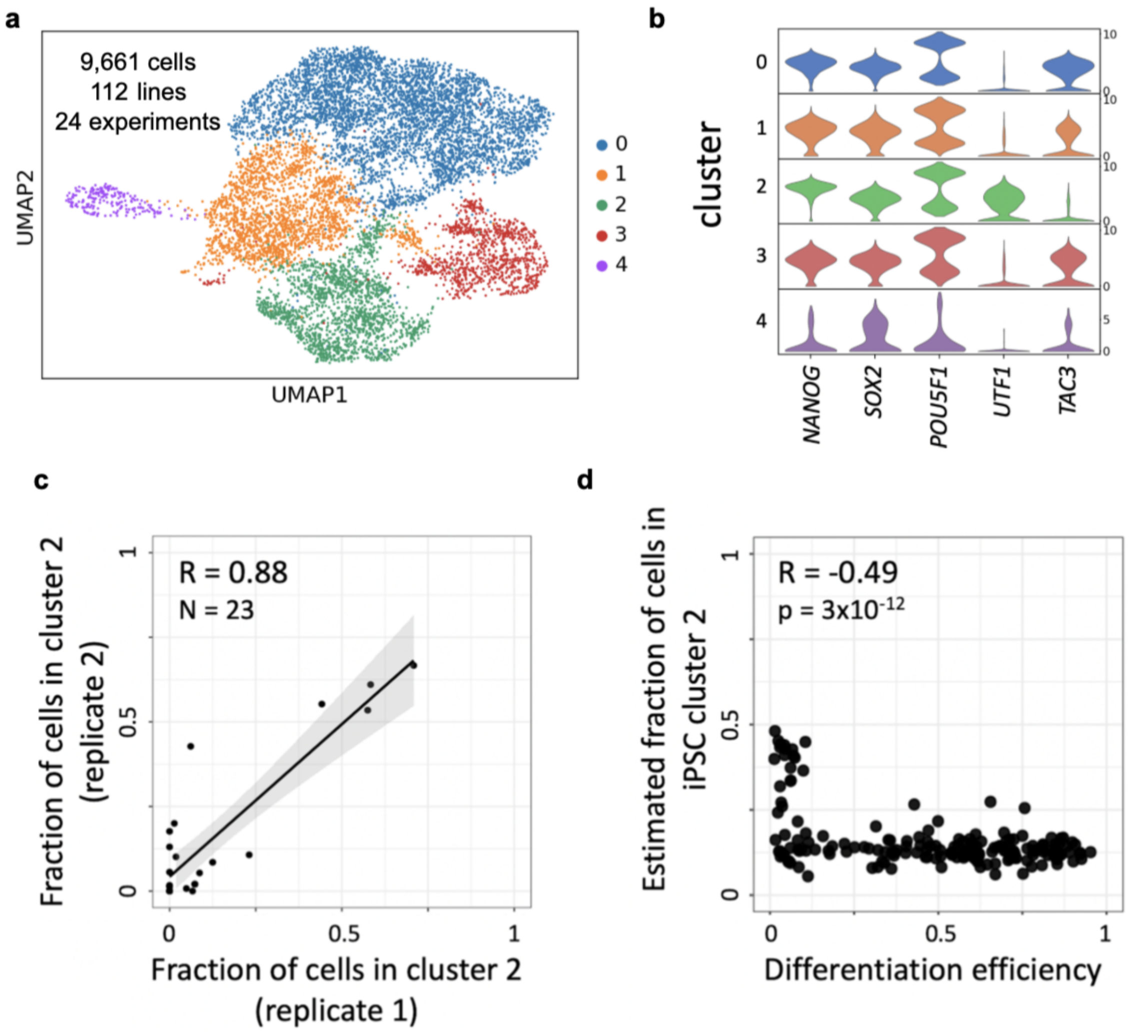

**Supplementary Figure 7.** Analysis of a single cell iPSC dataset from[4]. (a) UMAP overview of the dataset. iPSC scRNA-seq data from (Sarkar et al., 2019) were re-analysed following the same data normalisation and clustering steps applied to our neural differentiation data, identifying 4 clusters. (b) Violin plots of gene expression for *NANOG, SOX2, POU5F1*, *UTF1, TAC3,* from Fig. 3**,** Supplementary Figure 6. (c) Scatter plot showing the proportion of cells assigned to cluster 2 between replicates (n=59). (d) Expression log fold change between cluster 2 and all other clusters from [3] compared to the same between cluster 2 and the rest from [4]. Shown are all 5,397 DE genes between cluster 2 and all other clusters from Sarkar et al (FDR<0.05).

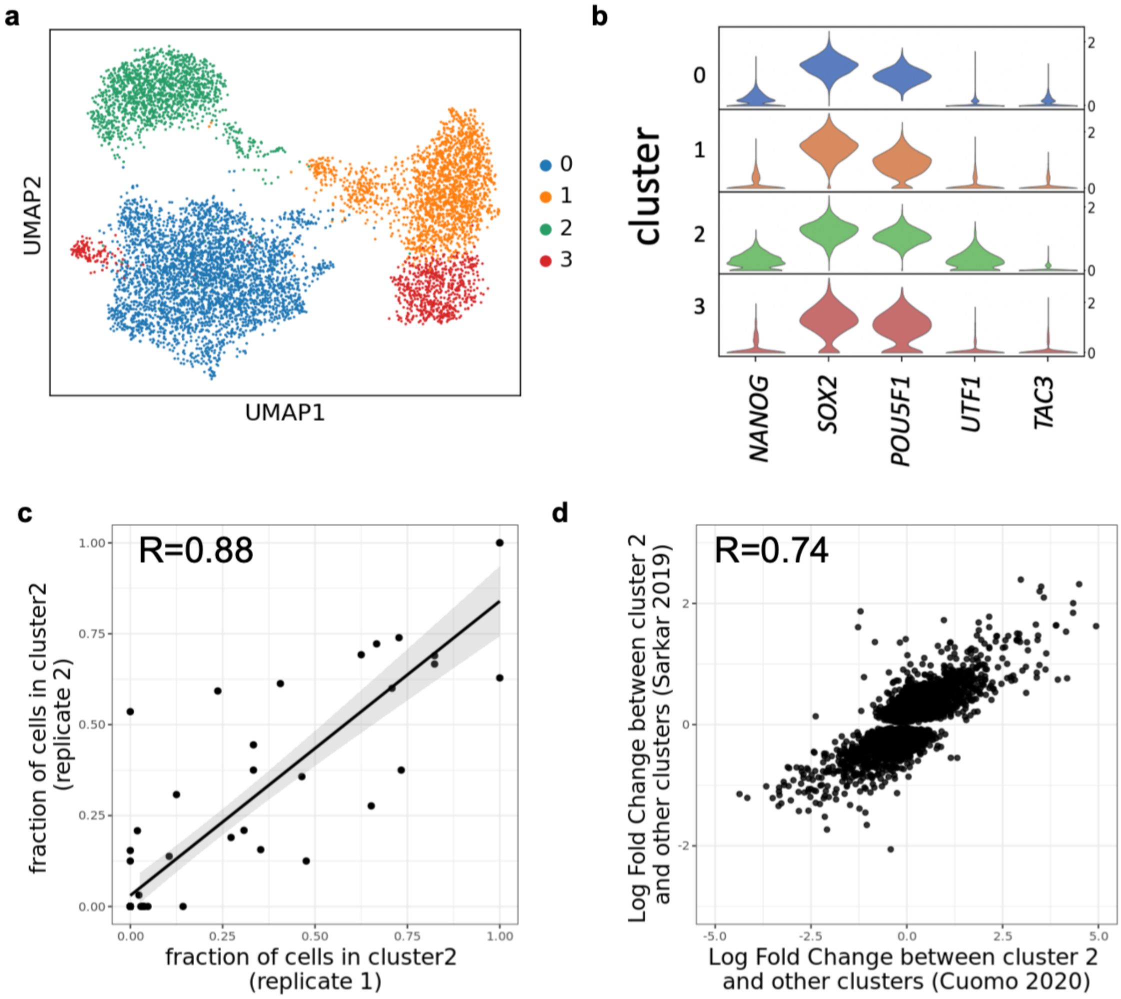

**Supplementary Figure 8.** Number of cells per cell type and condition and relation to eQTL power. (a) Distribution of the number of cells passing QC for each cell type and condition as obtained from 215 iPSC lines. (b) Number of genes with at least one eQTL (eGenes) for each cell type, and condition discovered using either a traditional linear model (coral) or our enhanced model that accounts for noise due to variation in the number of cells collected for each donor (seagreen). (c) Distribution of the number of cells passing QC for each cell type from 49 cell lines classified as good differentiators (differentiation efficiency>0.75). (d) Distribution of the number of cells passing QC **f**or each cell type and condition from the 52 cell lines classified as bad differentiators (differentiation efficiency<0.2 c.f. Supplementary Fig. 5a**).** Legend: Astro: Astrocytes-like; DA: Midbrain dopaminergic neurons, Epen1: Ependymal-like1, FPP: Floor Plate Progenitors, P_FPP: Proliferating Floor Plate Progenitors, Sert: Serotonergic neurons.

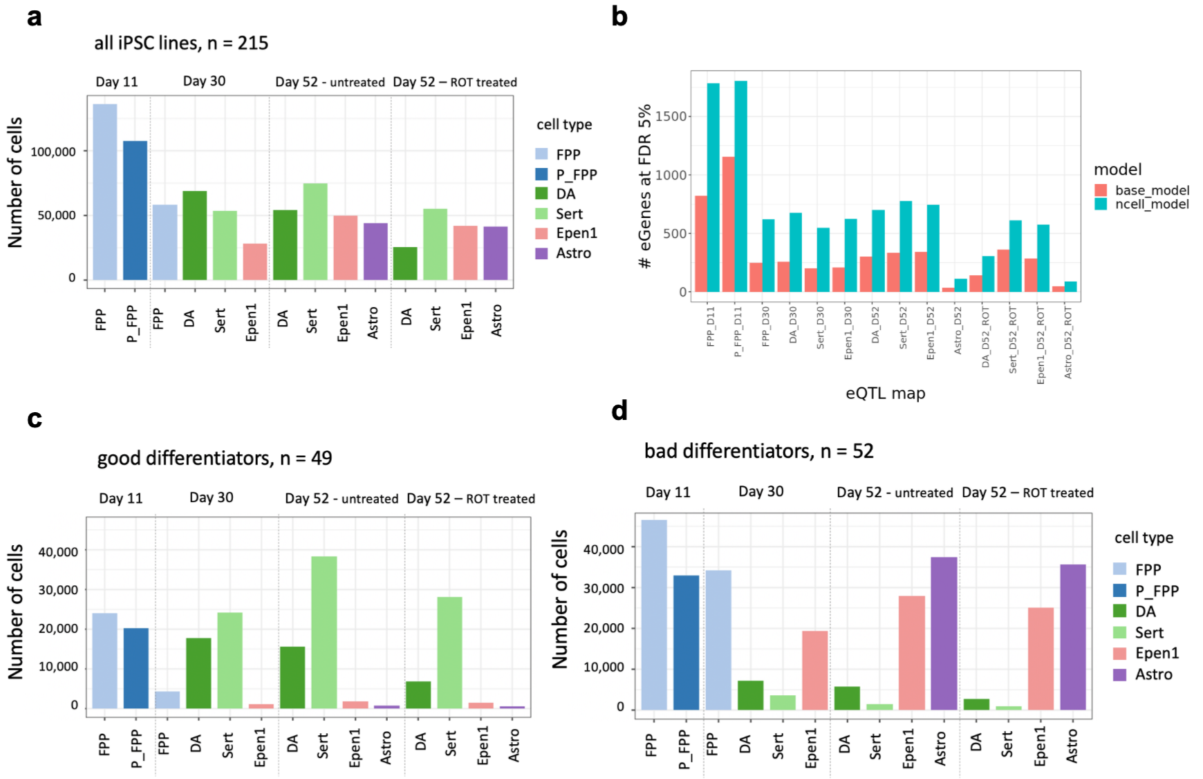

**Supplementary Figure 9:** eQTL and colocalisation. (a) Fraction of eGenes detected in each cell type, time point and stimulation that were not detected in each of the 13 brain tissue eQTL maps from GTEx (nominal p-value>0.05). (b) Similar to **Fig. 4a**. Cumulative number of colocalisation events with at least one neurological GWAS trait for each cell type (D11 = Day 11; D30 = Day 30; D52 = Day 52; ROT = Rotenone stimulation). Legend: Astro: Astrocytes-like; DA: Midbrain dopaminergic neurons, Epen1: Ependymal-like1, FPP: Floor Plate Progenitors, P_FPP: Proliferating Floor Plate Progenitors, Sert: Serotonergic neurons.

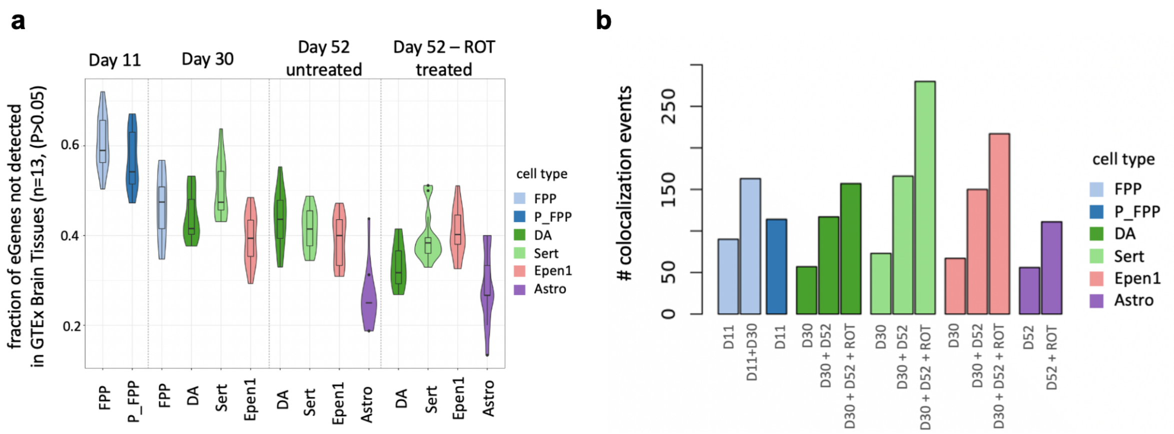

**Legend for Supplementary Tables:**
Note that all tables are supplied as external data files. For clarity, when fields are not readily interpretable we provide a small table to define them.

**Supplementary Table 1: Overview of the collected samples.**
All collected sample information including number of lines and replicated lines, single cell technical replicates and number of cells available per pool at each time point analysed are provided.

**Supplementary Table 2: Cell distribution per cell line, pool, cell type, time point and stimulation.**
Numbers of cells used in this study for each cell line per cell type, time point and stimulation.
Cell type: Astro: Astrocytes-like; DA: Midbrain dopaminergic neurons, Epen1: Ependymal-like1, Epen2: Ependymal-like2 FPP: Floor Plate Progenitors, NB: Neuroblasts, P_FPP: Proliferating Floor Plate Progenitors, Sert: Serotonergic neurons, U_Neur1: Unknown_neurons1, U_Neur1: Unknown_neurons2, U_Neur1: Unknown_neurons3

**Supplementary Table 3: Associations between cell lines' factors and differentiation efficiency, and between factors and predicted differentiation scores.**
Specified are the tests performed (Test), nominal p-values (p), sample size (n), and a True/False indicator of adjusted p-values (using Bonferroni) < 0.05. For continuous traits, Pearson correlation is shown (R), whilst for binary trait, the average difference is reported.

**Supplementary Table 4: List of genes correlated with differentiation efficiency**
Provides all identified significant positive and negative genes associated with differentiation efficiency at FDR <5%.

| **Table field** | **Description** |
|---|---|
| ensembl_gene_id | Ensembl ID (Ensembl version 75) |
| hgnc_symbol | Hugo Gene Nomenclature Committee symbol |
| coef | Effect size coefficient |
| pval | Nominal P-value |
| pval_adj | Adjusted P-value (Bejamini-Hochberg) |

**Supplementary Table 5: Predictive scores for HipSci banked lines**
Provides a predictive score of neuronal differentiation efficiency for the HipSci resource.
In the last column, cell lines that were included in this study were classified as "failed" (differentiation efficiency <= 0.2, n=48) or "succeeded" (differentiation efficiency > 0.2, n=136). Cell lines present in HipSci but not included in this study are "not-assessed" (n=628).

| **Table field** | **Description** |
|---|---|
| cell_line | HipSci cell line ID |
| model_score | Predicted score of differentiation efficiency (Methods) |
| diff_efficiency | Sum of the proportions of DA and Sert cells produced on day 52 |
| in_study | Cell line included or not in this study |

**Supplementary Table 6: DE genes between cluster 2 and others (single cell iPSC data).**
All differentially expressed genes between the cluster 2 and other clusters are provided after re-analysing iPSC scRNA-seq data from[3]. Results reported for FDR<5%.

| Table field | Description |
|---|---|
| ensembl_gene_id | Ensembl ID (Ensembl version 75) |
| hgnc_symbol | Hugo Gene Nomenclature Committee symbol |
| pval | Nominal P-value |
| pval_adj | Adjusted P-value (Benjamini-Hochberg) |
| log_fold_changes | Log Fold Changes |
| scores | Z-scores |
| cluster_id | Cluster ID |

**Supplementary Table 7: List of eQTL discoveries at FDR 5%**
All lead eQTL SNP-gene pairs are provided.

| Table field | Description |
|---|---|
| ensembl_gene_id | Ensembl ID (Ensembl version 75) |
| snp_id | Lead variant, SNP ID in the format [chromosome]_[position]_[reference]_[alternative allele] |
| label | Celltype and condition label, in the form [celltype]_[timepoint](_[stimulus]) |
| p_value | Nominal P-value |
| empirical_feature_p_value | Gene-level corrected P-value using using 1,000 permutations (Methods) |
| global_corr_p_value | Q-value, globally corrected P-value using Storey procedure (Methods) |
| beta | Effect size of the eQTL |
| beta_se | Standard error of the effect size |
| n_samples | Number of samples tested |
| feature_start | Gene body start position |
| feature_end | Gene body end position |
| snp_chromosome | Variant chromosome |
| snp_position | Variant position |
| assessed_allele | Variant allele assessed |
| maf | Variant minor allele frequency |
| hwe_p | Hardy-Weinberg equilibrium test p-value |

**Supplementary Table 8: List of neurological traits used for colocalisation analysis.**
All the neurological traits analysed are reported.

| Table field | Description |
|---|---|
| study_id | GWAS trait study ID (same as Supplementary Table 9) |
| pmid | PubMed unique identifier (when a publication is present and is not a preprint) |
| pub_date | Publication date |
| pub_journal | Publication journal (when a publication is present) |
| pub_title | Publication title (when a publication is present) |
| pub_author | First author of the publication |
| trait_reported | Trait reported |
| ancestry_initial | Ancestry of samples in main study |
| ancestry_replication | Ancestry of samples in replication study (when present) |
| n_initial | Sample size in main study |
| n_replication | Sample size in replication study (when present) |
| trait_category | Traits category |

**Supplementary Table 9: Results of the colocalisation analysis.**
All colocalisations identified are provided.

| Table field | Description |
|---|---|
| study_id | GWAS trait study ID (same as Supplementary Table 8) |
| celltype_tissue | Celltype/tissue label: [celltype]_[timepoint](_[stimulus]) for our study, tissue name for GTEx tissues |
| ensembl_gene_id | Ensembl ID (Ensembl version 75) |
| n_variants | Number of variants in 1Mb window |
| PP0 | Posterior probability 0 (no association with either trait) |
| PP1 | Posterior probability 1 (GWAS association, no eQTL) |
| PP2 | Posterior probability 2 (eQTL, no GWAS association) |
| PP3 | Posterior probability 3 (association for both, but independent SNPs) |
| PP4 | Posterior probability 4 (association for both, shared SNP) |
| chromosome_grch37 | Chromosome number according to GRch37 |
| GWAS_index_pos_grch37 | Position of in index GWAS variant 1Mb window |
| eQTL_lead_pos_grch37 | Position of lead eQTL in 1Mb window |
